# Mapping Uterine Remodeling from Pregnancy to a New Homeostatic State

**DOI:** 10.64898/2026.09.07.749876

**Authors:** Joji Marie Teves, Olga Wienskowska, Loris Mularoni, Borja Arozamena, Jordi Guiu

## Abstract

Pregnancy drives uterine remodeling through rapid anatomical expansion and scar-free postpartum repair, yet how the organ reconstructs its cellular architecture after birth remains incompletely understood. Here, we present single-cell transcriptomic profiling of 698,631 cells derived from the whole mouse uterus across five distinct stages centered around parturition: Non-pregnant Control, late gestation (embryonic day 16.5), and postpartum days (PPD) 1, 7, and 30. We resolved 16 distinct cell lineages spanning epithelial, stromal, endothelial, and immune compartments. Although gross morphological involution occurs rapidly, with uterine horn length returning to baseline levels by PPD7, the underlying cellular architecture does not revert to its pre-pregnancy state. Tissue composition, transcriptional states, and intercellular signalling networks did not revert to their pre-pregnancy configurations. Instead, postpartum reconstruction proceeded through five sequential but overlapping transcriptional programs, with individual cells simultaneously extinguishing gestational programs and acquiring repair-associated states. By one month postpartum, the tissue adopts a reconfigured architecture that fails to fully reactivate the non-pregnant baseline, thus, establishing a novel homeostatic state. This work provides a foundation for delineating the specific organizational levels that fully recover following physiological injury versus those that establish new baseline set points.

## Introduction

The endometrium exhibits a remarkable capacity for physiological tissue turnover that is unique among adult mammalian organs. Across the reproductive lifespan, it repeatedly dismantles and rebuilds its functional layer without undergoing fibrosis, structural degradation, or exhaustion of its regenerative potential (1–3). Whereas scarless tissue restoration is typically restricted to embryonic or neonatal stages, adult mammalian organs generally resolve injury through fibrotic scarring (4, 5). In contrast, the endometrium accomplishes scar-free reconstruction as a physiological routine and is therefore a tractable model to understand new insights about tissue regeneration (6).

Pregnancy presents a physiological challenge of even greater magnitude. Parturition detaches the placenta from the uterine wall, creating an acute, extensive wound at the implantation site. Subsequently, the organ must clear residual decidual tissue, degrade and rebuild the extracellular matrix, regress an expanded myometrium, and re-establish the luminal epithelium (7–9). Dysregulation of these postpartum repair processes can lead to significant clinical complications (10). The postpartum uterus is therefore a system in which cyclical physiological repair and large-scale post-injury reconstruction occur within the same resident cellular repertoire, providing a unique opportunity to investigate the extent and mechanisms of physiological regeneration.

Addressing this question has historically been limited by technical constraints. As repair proceeds simultaneously across epithelial, stromal, vascular, and immune compartments, sampling strategies restricted to isolated cell populations capture only part of the whole-organ process. Consequently, studies on endometrial progenitor populations have yielded conflicting conclusions regarding the cellular origins of the regenerated epithelium (11–14). For instance, the glandular epithelium has been proposed as a primary progenitor reservoir based on Foxa2 dependence and gland dynamics, although direct observation during postpartum repair remains limited (15–18). In contrast, a stromal contribution via mesenchymal-to-epithelial transition (MET) has also been reported in the postpartum mouse model (19, 20). Cross-sectional analyses are further confounded by estrous cycle fluctuations, which drive substantial transcriptional, compositional, and structural variations in the rodent uterus independently of tissue injury (21, 22).

Single-cell transcriptomics has advanced our understanding of female reproductive tract biology, yet existing datasets have largely omitted the postpartum window. Available single-cell atlases cover the cycling and aging mouse reproductive tract (23), cycling human endometrium and its chromatin dynamics (24–27), healthy premenopausal human uterus (28), the mouse endometrial mesenchymal compartment (29), maternal-fetal interface throughout gestation (30–32), and menstrual or disease-associated endometrium (33–35). Studies focused on peripartum dynamics have primarily captured the term human myometrium or murine labor models rather than the subsequent involution phase (36–38). Consequently, uterine involution has been characterized mainly through histology, targeted molecular assays, and bulk transcriptomics (8, 9, 39). A whole-organ single-cell map of postpartum repair anchored at parturition has been lacking, leaving a central question unanswered: whether histological recovery during involution reflects a genuine return to the pre-pregnancy state.

To address this gap, we performed single-cell RNA sequencing on the whole dissociated uteri, capturing all major tissue compartments simultaneously to enable an integrated analysis of cross-compartment signaling. We profiled 698,631 cells across 15 libraries, 14 of which passed quality control, spanning non-pregnant control, late gestation (E16.5), and postpartum days 1, 7, and 30, resolving 16 distinct cell types. Including multiple postpartum time points allowed us to segregate the acute response to parturition from long-term tissue remodeling. To directly link gross tissue dynamics with molecular changes, we paired single-cell transcriptomics with anatomical measurements of uterine horn length and EPCAM/FOXA2 immunofluorescence on contralateral uterine horns from the same animals.

Our analyses reveal that postpartum recovery is stratified across biological scales. Gross anatomy returns to baseline within one week, yet this apparent restoration conceals persistent reorganization of cellular composition, transcriptional states, and intercellular signalling networks. We resolved this process into five sequential transcriptional programs that display temporal overlap within individual cells rather than in distinct cell subpopulations. These shared programs unfold asynchronously across tissue compartments, with the immune compartment following distinct temporal dynamics. In parallel, the intercellular communication network is progressively rewired and remains markedly displaced from the non-pregnant configuration at one month postpartum. Thus, the postpartum uterus does not simply reverse pregnancy-induced changes, instead it restores organ structure while establishing a distinct cellular and molecular configuration.

## Results

### Postpartum uterine involution attains anatomical baseline within one week

To benchmark cellular and molecular dynamics against physiological organ restoration, we first delineated the anatomical trajectory by measuring the gross uterine expansion (Fig. 1a). Uterine horn length expanded more than threefold during gestation, peaking at 55.3 mm at E16.5, before undergoing rapid postpartum involution to 26.8 mm by PPD1 and 21.1 mm by PPD7, against a non-pregnant Control value of 16.5 mm (Fig. 1b). Gross anatomical restoration was completed within one week of parturition (Fig. 1c). Representative whole-tissue images corroborate this rapid morphological trajectory (Fig. 1d). Together, these findings confirm that macro-scale morphological deviations relative to the pre-pregnancy organ resolve within the first week postpartum.

**Fig. 1.**
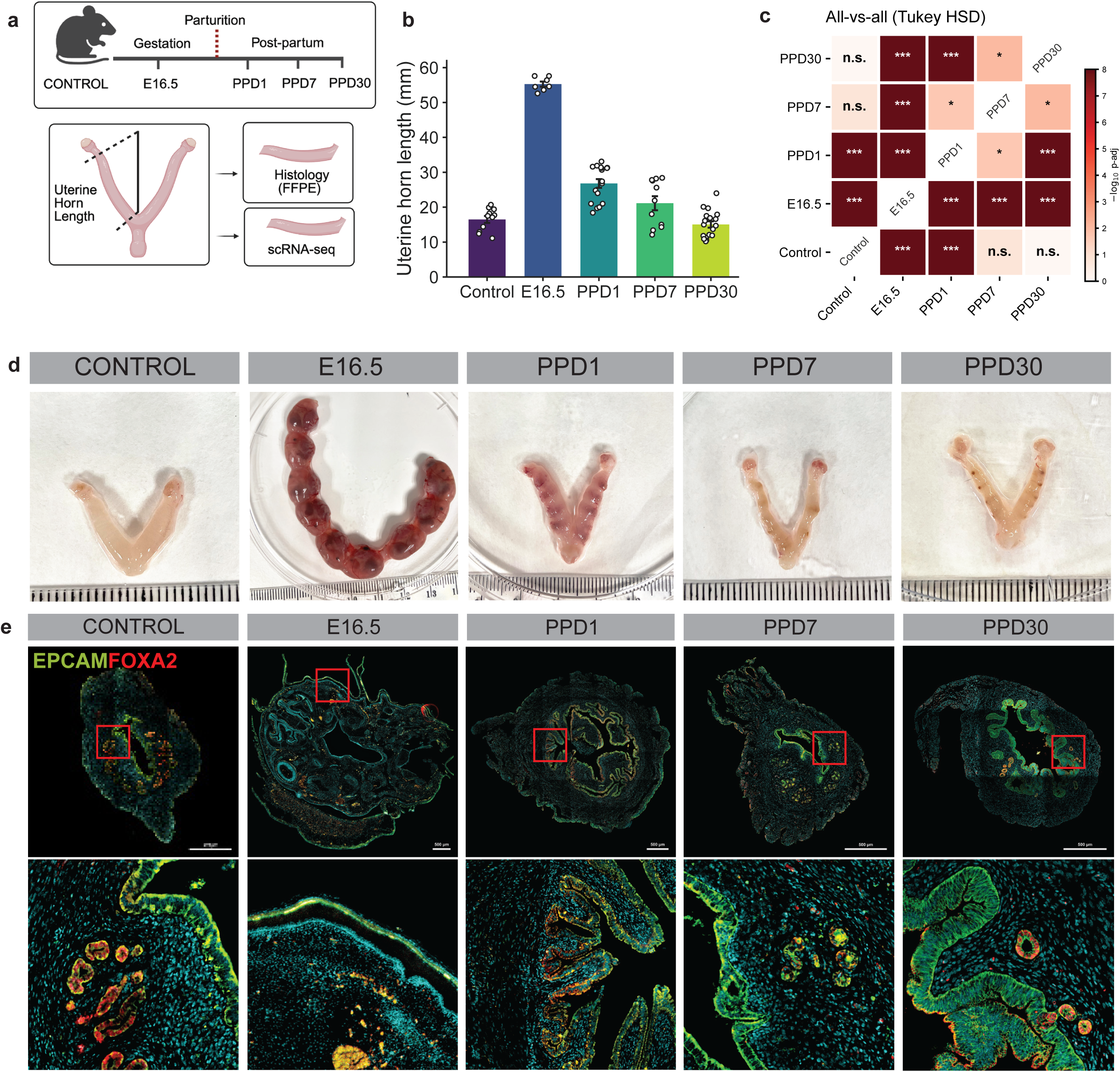
Gross involution of the postpartum uterus completes within one week and is sustained through three months, whereas luminal epithelial continuity is re-established over one month. (**a**) Experimental design. Whole uteri were collected from non-pregnant control adult females (CONTROL), at embryonic day 16.5 of gestation (E16.5), and at postpartum days 1, 7 and 30 (PPD1, PPD7, PPD30), with parturition defining time zero. One uterine horn was fixed for histology and the contralateral horn dissociated for single-cell profiling, such that anatomical and transcriptional measurements are paired within animal. Animals collected at postpartum day 90 (PPD90) contributed anatomical measurement only. (b) Uterine horn length by stage: CONTROL 16.5 mm (3.3, 11), E16.5 55.3 (2.0, 7), PPD1 26.8 (5.2, 17), PPD7 21.1 (6.7, 11), PPD30 15.0 (3.9, 18), PPD90 12.1 (1.4, 4). (c) All-versus-all comparison of horn length across groups by one-way ANOVA with Tukey HSD, where cell color encodes –log10 adjusted P with signifcance classification (n.s., *P* ≥ 0.05; * ! *P* < 0.05; **, *P* < 0.01; ***, *P* < 0.001) (d) Representative whole-tissue images of uterine horns at six stages with ruler graduations, 1 mm. (e) Cross-sections stained for EPCAM (green) and FOXA2 (red) with nuclear counterstain (blue), for five stages; Upper row, whole uterus cross-section, with the red box indicating the region magnified in the lower row; lower row, magnified view. Scale bars, *[500µm]*.

In contrast to the rapid recovery of organ-level dimensions, epithelial reconstruction follows a markedly prolonged trajectory. Immunofluorescence profiling for EPCAM and FOXA2 revealed an attenuated, discontinuous luminal epithelium at PPD1 and PPD7, demonstrating compromised tissue integrity during early repair. By PPD30, however, a continuous EPCAM-positive luminal layer accompanied by fully formed FOXA2-positive endometrial glands is restored (Fig. 1e). Thus, organ size and epithelial continuity recovered on distinct timescales. We therefore selected stages spanning non-pregnant control, late pregnancy (E16.5), acute postpartum injury (PPD1), ongoing repair (PPD7), and restored epithelial homeostasis (PPD30) for whole-uterus single-cell profiling.

### The cell-type repertoire is preserved while compartment composition is extensively remodelled

To map the cellular architecture across postpartum tissue restoration, we dissociated the whole uterus to single cells at five stages. To ensure adequate recovery of viable cells for single-cell capture, we optimized the collagenase digestion time-course (30, 45 and 60 mins; Fig. S1a-f).

We identified 45 mins as optimum for cell yield and compartment recovery. This approach recovered 698,631 single cells of which 698,595 lie in the fourteen libraries retained for replicate-level analysis after quality control (Table S1, Fig S1g-l). Importantly, equal-N subsampling reproduced the direction of every compositional change (Fig. S1h),to ensure that unequal cell recovery across stages do not confound compositional shifts. Batch-corrected integration of 15single-cell RNA-sequencing libraries yielded a single embedding in which biological replicates and stages intermix without batch-driven segregation (Fig. 2a, b). Unsupervised clustering combined with canonical marker gene annotation resolved 16 cell types spanning the epithelial, stromal, endothelial, and immune compartments (Fig. 2b, c). Each distinct cellular identity was validated by a tailored transcriptional marker program defined across 139 genes (Fig. 2d). Sensitivity analyses confirmed that lineage annotations were stable across multiple clustering resolutions and were independent of technical confounders, including sequencing depth, transcript counts, mitochondrial read fractions, and predicted doublet scores (Fig. S2a-c). All 16 cell types were detected at every stage, so the non-pregnant control repertoire is preserved in full across parturition and repair (Fig. 2e). This conservation of lineage identity indicates that the dynamic tissue-level transformations observed throughout involution reflect profound shifts in cellular state and proportion rather than the complete deletion or de novo emergence of specific cell types.

**Fig. 2.**
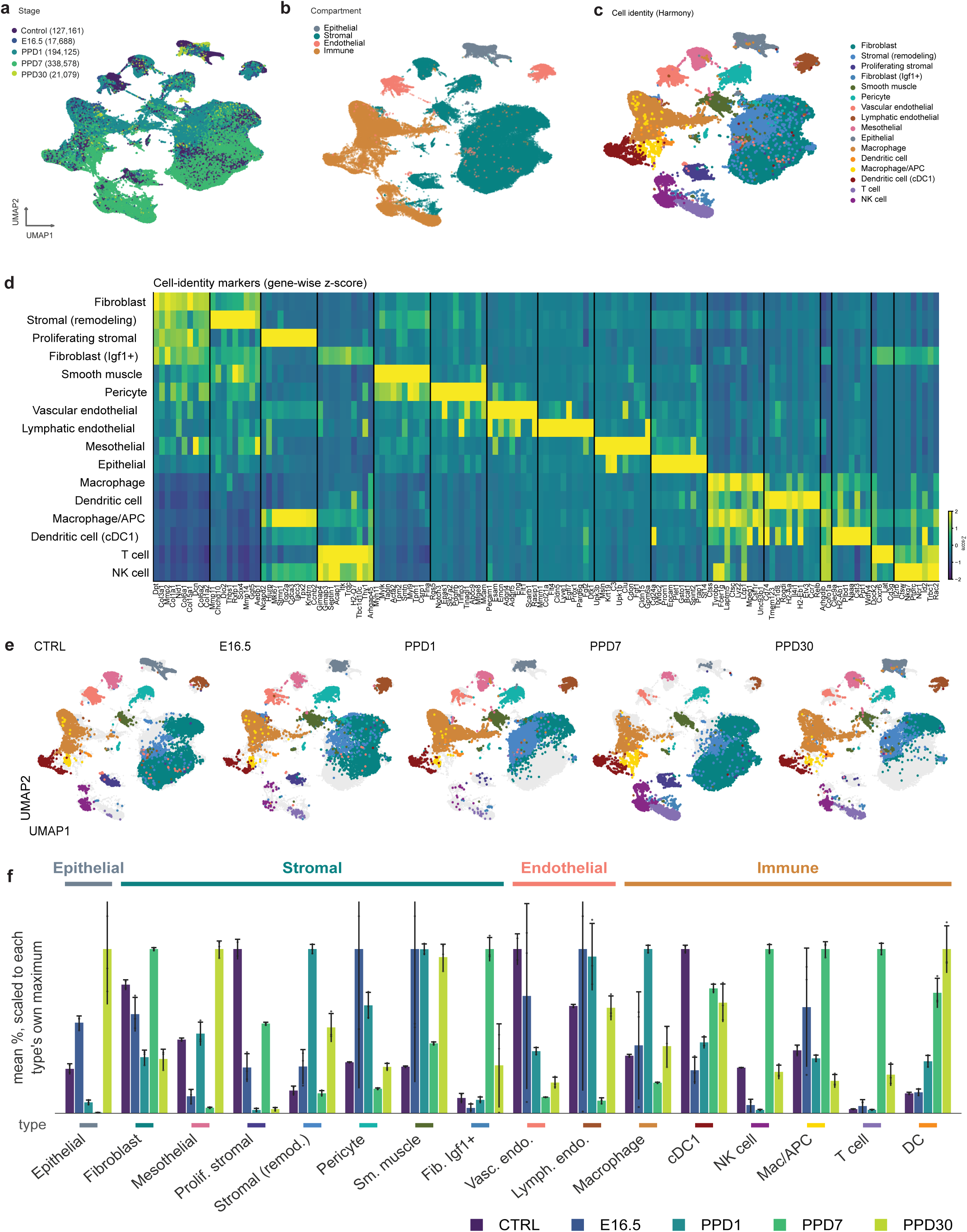
A whole-uterus single-cell reference spanning parturition, in which the cellular repertoire is preserved while compartment composition is extensively remodeled. (**a**) Uniform manifold approximation and projection (UMAP) of 698,631 Harmony-integrated cells colored by stage. (b, c) The same embedding colored by compartment (b) and by cell type (c). (**d**) Gene-wise, standardized expression of 139 cell-identity markers across the 16 cell types, in which each gene is *z*-scored across types. (**e**) Stage-resolved projections, with cells of the indicated stage in colour and all remaining cells in grey where every cell type is detected at every stage. (**f**) Mean per-replicate abundance of each cell type, scaled to that type’s maximum across stages, with bars showing mean and whiskers showing SD across *2–3 sequencing* libraries per stage; absolute proportions are provided in Table S2.

In contrast, cellular composition shifted markedly across stages (Fig. 2f, Fig. S2d). The epithelial-to-stromal ratio (Fig. 2f), measured at 0.101 in non-pregnant control, rose to 0.223 during late gestation (E16.5) before declining sharply to 0.028 at PPD1 and 0.002 at PPD7. By PPD30, this ratio increased to 0.485, representing a nearly fivefold expansion relative to the non-pregnant control baseline. Across these stages the embedding contains 127,161 cells from the non-pregnant controls, 17,688 at E16.5, 194,125 at PPD1, 338,578 at PPD7, and 21,079 at PPD30 (Fig. 2e); the corresponding retained-library counts are given in Table S1. We further compared our annotation to a published whole-uterus reference, correlating compartment-level pseudobulk profiles against the Mouse Cell Atlas (69), and found that each of our six compartments correlated most strongly with its own counterpart in the reference (diagonal Pearson r 0.49–0.73, the row maximum in every case; Fig. S2e-f). The distinct temporal peaks in relative abundance observed across specific cell lineages suggest that tissue compartments undergo remodeling according to asynchronous timescales.

Moreover, transcriptional output varied across the remodeling trajectory, reaching its maximum at E16.5, with 8,285 mean unique molecular identifiers per cell compared with 2,279 in non-pregnant controls (Table S1), consistent with the extensive cellular activation accompanying pregnancy. Putatively, the compositional shifts reflect broad biological reorganization of the uterus across pregnancy and postpartum repair rather than differences in sequencing representation or overall tissue viability. Collectively, these findings demonstrate that although lineage identities are maintained, their relative proportions undergo extensive structural reorganization.

### Five transcriptional programs peak in succession, and the Homeostatic program is never re-established

To directly characterize cell-state trajectories, we derived gene modules from whole-tissue pseudobulk differential expression analysis against the non-pregnant control and evaluated module scores across all individual cells (Fig. S3a). This approach identified five distinct transcriptional programs, each attaining maximum expression at a successive stage: Homeostasis (678 genes, maximal in control), Pregnancy (217 genes, E16.5), Early repair (985 genes, PPD1), Late repair (745 genes, PPD7), and New Homeostasis (789 genes, PPD30) (Fig. 3a; Table S3). The programs followed overlapping temporal waves with each module progressively rising as the previous modules declined (Fig. 3b–f, Fig S3b). Programs were scored on the stable 16-type annotation (43, 44) and robustness metrics for these programs were assessed against size-and expression-matched random gene sets, per-cell-type trajectories, and pairwise program overlaps(Fig. S3c and Table S3).

**Fig. 3.**
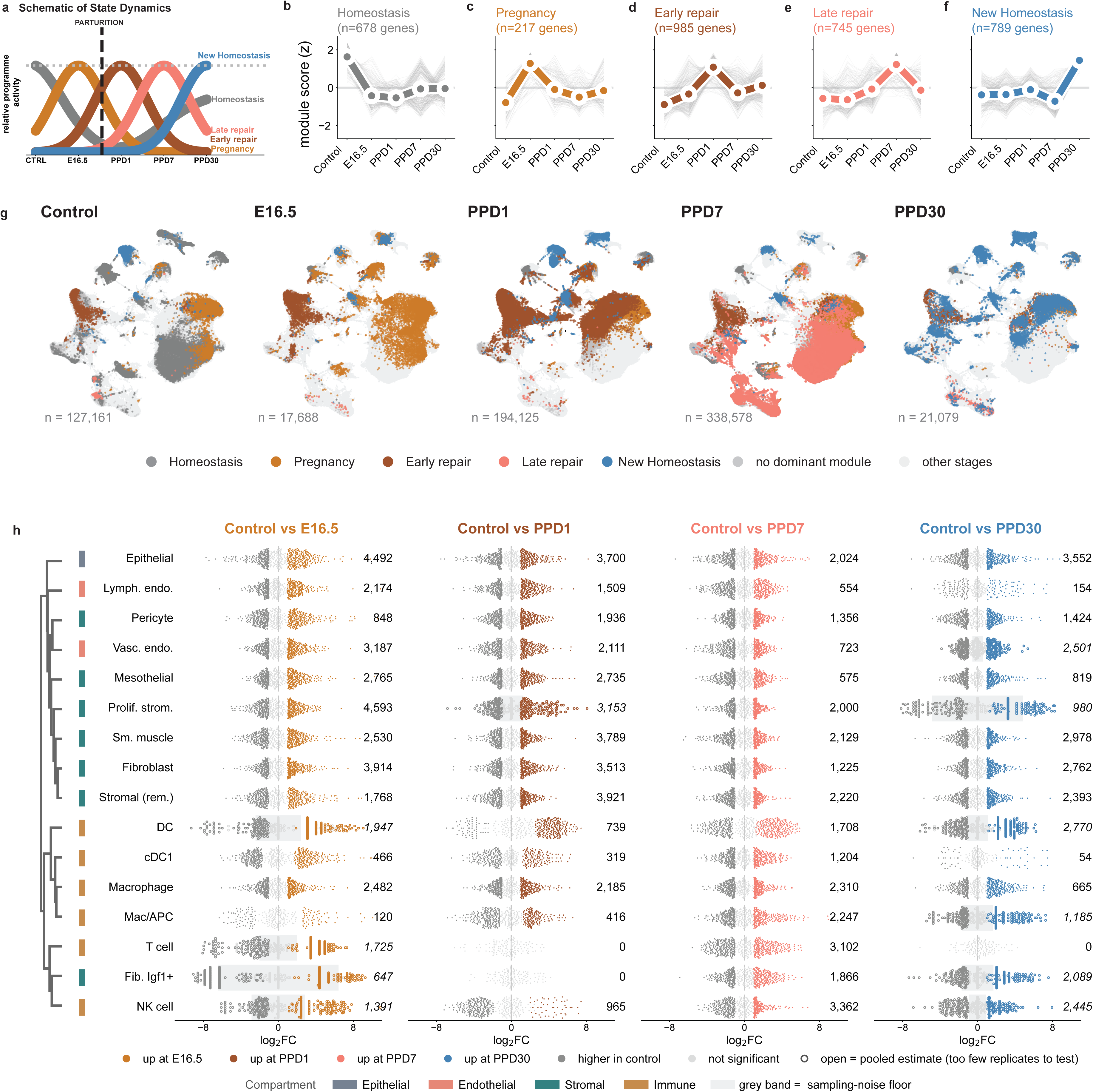
Five transcriptional programs attain their maxima sequentially across parturition, and the Non-pregnant control program is not re-established at any postpartum stage. (**a**) Schematic of the sequential program hand-over inferred from **b–f**, with parturition indicated. (**b– f**) Module scores across stages for Homeostasis (678 genes), Pregnancy (217), Early repair (985), Late repair (745) and New Homeostasis (789), in which grey traces denote individual genes and the colored trace the module mean. (**g**) Dominant program per cell projected onto the UMAP embedding. (**h**) Differential expression relative to CONTROL for each cell type at each stage.

The baseline Homeostasis program failed to fully recover after parturition. Its expression score remained significantly below non-pregnant control levels at all postpartum time points, including PPD30, a stage at which gross anatomical parameters had been indistinguishable from baseline for over three weeks (Fig. 3b). Dominant-program assignment at single-cell resolution revealed a sequential transition from pregnancy through early and late repair, culminating in a New Homeostasis at PPD30 (Fig. 3g). Postpartum recovery therefore does not retrace the trajectory of pregnancy in reverse but instead, repair resolves into a new transcriptional state rather than restoring the original homeostatic program.

Additionally, this transcriptional displacement was not restricted to the acute phase and exhibited its greatest magnitude within the epithelial compartment. Cell-type-specific pseudobulk differential expression placed the epithelium among the most transcriptionally displaced types at every stage, and made it the single most displaced type at PPD30: 4,492 genes at E16.5, 3,700 at PPD1, 2,024 at PPD7, and 3,552 at PPD30 (Fig. 3h). Transcriptional divergence at one month postpartum thus approached that of late gestation. To avoid false-discovery inflation caused by treating single cells as independent observations, differential expression testing was performed on aggregated pseudobulk profiles per replicate and cell type (45–48).

Unlike the gestational transcriptional trajectory, the immune compartment exhibits distinct dynamics. Aggregating significantly altered genes across all six immune cell types yields 8,131 genes at E16.5, 4,624 at PPD1, 13,933 at PPD7, and 7,119 at PPD30 (Fig. 3h). Whereas the whole-tissue Early repair program peaks at PPD1, immune divergence falls between gestation and PPD1 and reaches its maximum at PPD7. Individual immune lineages show the same offset. The Macrophage/APC lineage, for example, rises from 120 differentially expressed genes at E16.5 to 416 at PPD1 and 2,247 at PPD7 (Fig. 3h). Immune involvement thus peaks during late repair rather than during late gestation or the acute postpartum phase, providing initial evidence of inter-compartmental desynchronization.

Overall, the uterus had not recovered its pre-pregnancy transcriptional state by one month postpartum. However, whole-tissue programs cannot determine whether this persistent displacement is shared across compartments or driven by specific cell lineages.

### Each compartment executes the shared programs on independent timescales

To resolve compartment-specific dynamics, we scored each transcriptional program across individual cell lineages and the four major tissue compartments (Fig. S3d-e). All five programs were detectable across compartments, but their temporal trajectories differed (Fig. 4a–e). Four programs reached maximal expression at their defining stages in all four compartments. The Late repair program showed a distinct epithelial trajectory, with lower overall activity, an earlier peak at E16.5, and sustained elevation through PPD7. By contrast, stromal, endothelial, and immune compartments exhibited a more temporally restricted response that peaked at PPD7. Thus, although the same transcriptional programs operate across the uterus, each compartment executes them with distinct timing and amplitude.

**Fig. 4.**
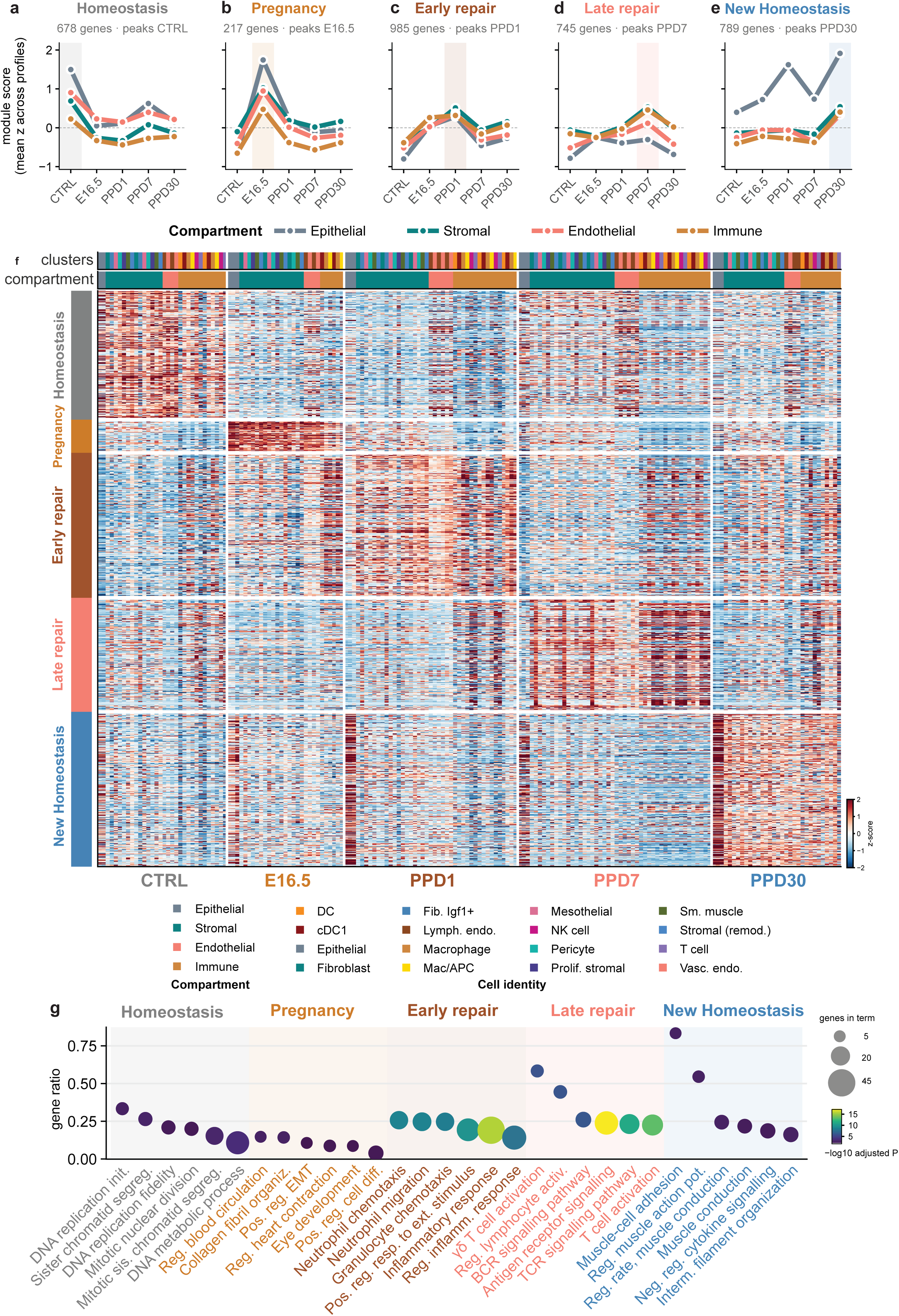
Each compartment executes the shared transcriptional programs on an independent schedule. (**a**–**e**) Module score per compartment across stages, one panel per program, with the program’s peak stage shaded; points denote the mean standardized score across replicate and cell-type profiles within compartment. (**f**) Program-defining genes as rows, grouped by program with the program colour bar at left, across replicate and cell-type pseudobulk profiles as columns, grouped by stage, with annotation tracks for cell identity and compartment above; values are gene-wise *z*-scores across profiles. (**g**) Gene Ontology biological process terms enriched wihtin each program

Notably, the epithelium exhibited a pronounced, non-monotonic expression trajectory for the New Homeostasis program, rising at PPD1, dropping at PPD7, and reaching a maximum at PPD30 that substantially exceeded levels in other compartments (Fig. 4e). Consequently, the epithelial compartment undergoes the most extensive structural depletion during early repair and displays the greatest transcriptomic divergence from the non-pregnant control baseline at one month postpartum. Rather than reinstating its pre-pregnancy program, the restored epithelium acquired a new homeostatic state that retained a transcriptional imprint of the preceding pregnancy-to-repair transition.

These compartment-level dynamics were consistently preserved across individual replicate-by-cell-type profiles, confirming that the observed patterns do not arise from sample aggregation artifacts (Fig. 4f). At cell-type resolution, program activation aligned with the expected peak stage in 16 of 16 cell types for Late repair, 13 of 16 for Early repair, 13 of 16 for New Homeostasis, and 11 of 16 for Pregnancy, with every dissenting lineage belonging to the immune compartment (Fig. 4f). The Homeostasis program remained suppressed below control across all 13 sufficiently covered cell types at PPD30 (Fig. 4f), confirming that the failure to reactivate the pre-pregnancy baseline program is a tissue-wide phenomenon. Functional annotation follows an ordered sequence of repair processes (Fig. 4g). The Pregnancy program enriched for regulation of vascular circulation, extracellular matrix organization, and positive regulation of epithelial-to-mesenchymal transition. Early repair enriched for innate immune activation, including neutrophil and granulocyte chemotaxis. Late repair was dominated by adaptive immune processes, including T cell and B cell receptor signaling, so adaptive recruitment follows the initial inflammatory phase. New Homeostasis enriched for muscle-cell adhesion, regulation of muscle action potential, intermediate filament organization, and negative regulation of cytokine signaling, reflecting myometrial stabilization and the active resolution of inflammation. In contrast, the Homeostasis program was predominantly enriched for DNA replication, mitotic nuclear division, and chromosome segregation. This indicates that the non-pregnant control homeostatic signature reflects a baseline proliferative state. Thus, postpartum reconstruction follows a common transcriptional sequence, but its temporal execution is putatively compartment-specific, generating distinct functional trajectories within the same organ-wide transition. We therefore next resolved how each compartment contributes to the New Homeostasis state and identified the biological processes associated with its distinct transcriptional configuration.

Compartment-resolved analysis revealed that New Homeostasis is not a uniform organ-wide program, but a composite state built from distinct transcriptional and functional responses in the epithelial, stromal, endothelial, and immune compartments (Fig. 5a–d). Gene Ontology enrichment further showed that each compartment contributed a different biological component to this reconfigured state (Fig. 5e–h). This compartment specificity was most pronounced in the epithelium, where Inflammatory Response (GO:0006954) was the single most enriched term of the epithelial New Homeostasis module, covering 16 of its 313 genes (gene ratio 0.051; adjusted P = 6.4 × 10⁻³), alongside Regulation of Wound Healing (7 genes; adjusted P = 6.4 × 10⁻³). The contributing genes include the interleukin-1 family members Il1a, Il36a, Il36b and Il36g, the innate sensors Nlrc4, Zbp1 and Tlr5, and the NF-κB subunit Relb, indicating that the epithelium carries a constitutively upregulated inflammatory program into its new homeostatic state. This signature is epithelium-specific wherein the same term ranks eighteenth in the immune module (adjusted P = 0.056) and is not enriched in the stromal or endothelial modules (adjusted P = 0.74 and 0.89). While individual compartments execute shared programs along distinct temporal schedules, inter-compartmental coordination cannot be driven solely by intrinsic cell-autonomous programs. Thus, the desynchronization suggests that tissue restoration relies heavily on extrinsic cell-cell communication networks to orchestrate cross-compartment repair.

**Fig. 5.**
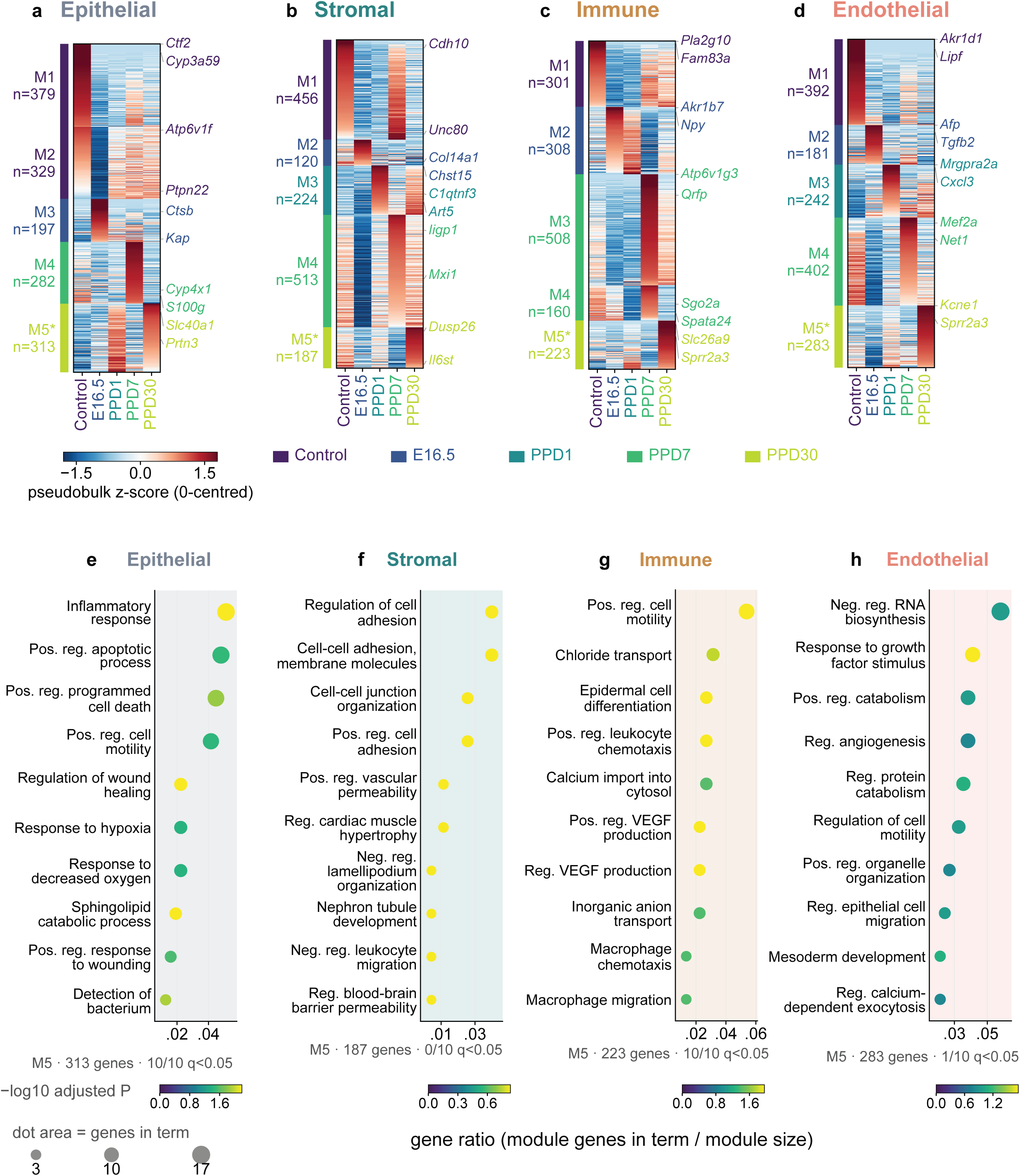
Each compartment builds the New Homeostasis state from a different transcriptional module. (**a-d**) Per-compartment pseudobulk profiles were computed across the five stages; the 1,500 genes of highest one-way ANOVA F across stages were retained per compartment and partitioned into five temporal modules (M1-M5) by k-means on stage z scores. Module structure for the epithelial (**a**), stromal (**b**), immune (**c**) and endothelial (**d**) compartments(**e-h**) Gene Ontology biological-process enrichment of each compartment’s M5 module, run independently per compartment against an all-genes background with Benjamini-Hochberg correction, for the epithelial (**e**), stromal (**f)**, immune (**g**) and endothelial (**h**) compartments.

### The intercellular communication network is progressively rewired and does not revert

Given the desynchronized transcriptional dynamics observed across tissue compartments during orderly repair, we reasoned that intercellular signaling networks must orchestrate this compartmental coordination. Utilizing whole-organ single-cell profiling, ligand-receptor communication networks were inferred at each stage to evaluate whether intercellular signaling channels revert to the non-pregnant control homeostatic baseline.

We observed that signalling roles shifted across stages within every compartment (Fig. 6a–d, Fig. S4a-c). The dominant stromal sender shifted from mesothelial cells at E16.5 (5,825 interactions) to fibroblasts at PPD30 (5,480 interactions). Meanwhile, epithelial sender activity expanded progressively from E16.5 through PPD30, whereas lymphatic endothelial cells remained the primary endothelial sender at every stage. Within the immune compartment, macrophage populations carried the highest signalling output at E16.5. Network role assignments thus remain dynamic despite the conservation of lineage identities across time. Compositionally, the signaling network did not converge toward the non-pregnant control state (Fig. 6e) wherein every stage gained more interaction edges than lost. Jaccard similarity to the control network peaked at 0.431 at PPD7, the most baseline-like postpartum stage, before declining to 0.347 at PPD30 (Fig. 6e). Consequently, the network transiently approached the non-pregnant control baseline at one week postpartum before undergoing secondary divergence. Total inferred interactions numbered 29,108 in controls, 49,109 at E16.5, 42,870 at PPD1, 40,967 at PPD7, and 53,561 at PPD30 (Fig. 6e). Intercellular interactions thus reached a postpartum minimum at PPD7 (remaining 1.41-fold above baseline) and peaked at PPD30, surpassing even late-gestational totals. Moreover, net Jaccard set similarity provides the primary basis for network assessment, as edge-set similarity metrics are less susceptible to sequencing depth variations across stages than raw interaction counts.

**Fig. 6.**
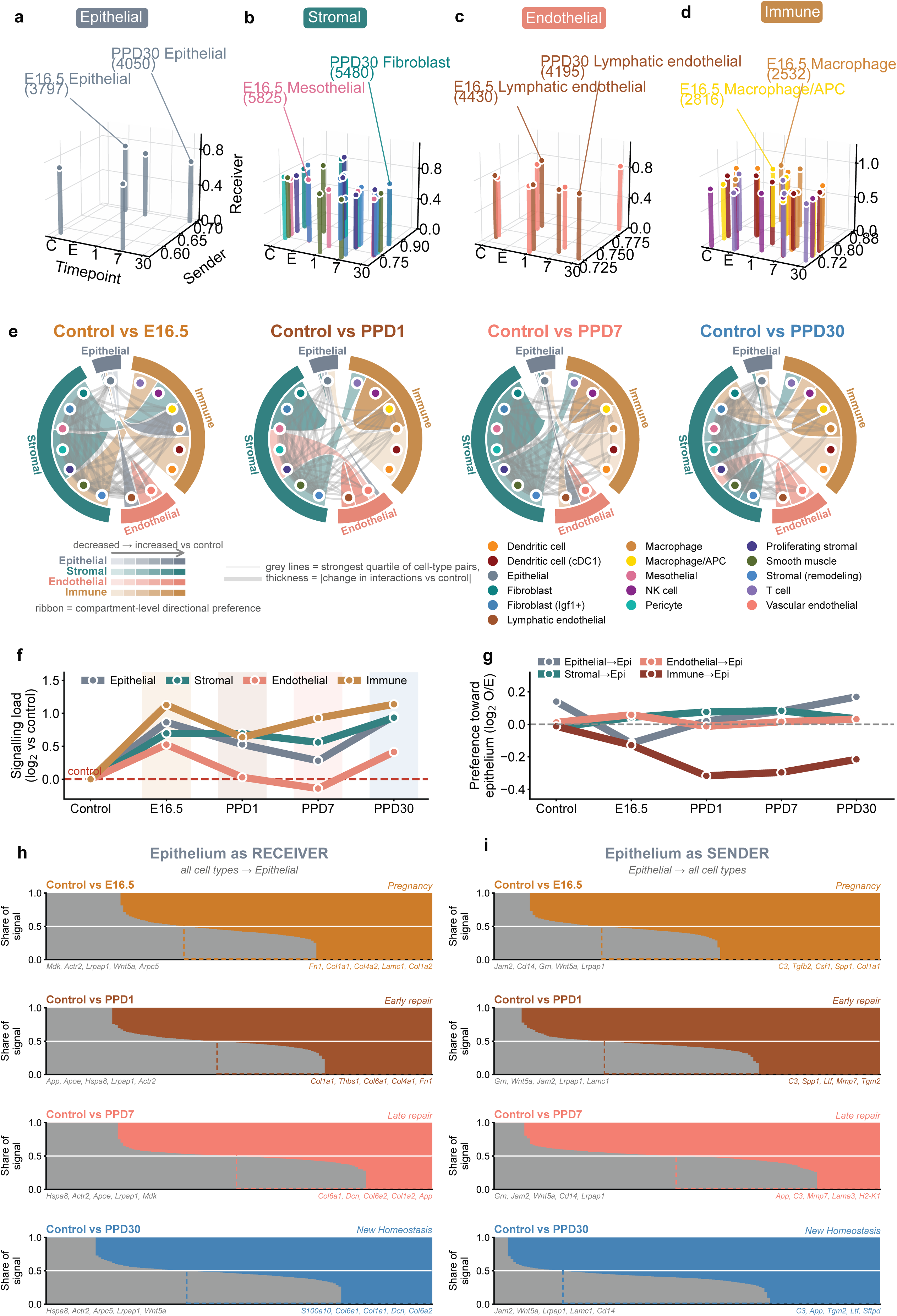
Inferred signalling load is elevated in every compartment during New Homeostasis (PPD30). Ligand–receptor interactions were inferred across all 16 annotated cell types and every comparison is anchored on the non-pregnant control. (**a-d**) Sender and receiver activity per cell type, one panel per compartment. Each stem marks one cell type at one timepoint, positioned by its mean interaction score as a sender (depth axis) and as a receiver (height axis); color identifies the cell type. Leader lines name the two cell types with the largest number of significant outgoing interactions in that compartment across all stages. (**a**) epithelium 4,050 at PPD30 and 3,797 at E16.5; (**b**) mesothelial 5,825 at E16.5 and fibroblast 5,480 at PPD30; (**c**) lymphatic endothelial 4,430 at E16.5 and 4,195 at PPD30; (**d**) macrophage/APC 2,816 and macrophage 2,532, both at E16.5. (**e**) Control-anchored network change with one disc per postpartum comparison. Ribbons carry compartment-level change in directional preference, with lightness encoding direction. Grey lines are individual cell-type pairs with thickness proportional to the absolute change in interaction count; (**f**) Total signalling load per compartment, log₂ relative to control. (**g**) Directional preference toward the epithelium for each sender compartment. (**h**) Epithelium as receiver with every cell type signalling to the epithelium. (**i**) Epithelium as sender: the epithelium signalling to every cell type.

Nevertheless, total signaling load remained elevated across all tissue compartments at one month postpartum (Fig. 6f). At PPD30, the immune compartment carried the greatest signaling load relative to control (1.14 log₂), followed by the epithelial (0.96), stromal (0.93) and endothelial (0.41) compartments (Fig. 6f), with each reaching or nearing its maximum activity level. The endothelial compartment was the only tissue domain to fall below baseline signaling levels at any stage, doing so transiently at PPD7. Directional targeting analyses established that the restored epithelium occupies a signaling position distinct from that of the non-pregnant control epithelium (Fig. 6g). Epithelial self-preference recovered to 0.169 at PPD30, exceeding the non-pregnant control baseline of 0.140 (Fig. 6g, Fig S4c-d). This is accompanied by changes in the signals received and sent by the epithelium across the different stages (Fig, 6h-i). Conversely, immune-to-epithelial signaling preference became strongly negative at PPD1 (-0.317) and remained persistently suppressed at PPD30 (-0.216), whereas the stromal-to-epithelial preference remained positive throughout. Thus, at one month postpartum, the restored epithelium interacts predominantly with itself while the immune inputs remain directed away from it which represent a configuration opposite to an expected from a return to the pre-pregnancy state. Thus, these transcriptional modules reflect distinct biological programs wherein the modules were derived from pseudobulk differential expression analysis, and the intercellular interactions were inferred from single-cell co-expression profiles. Genes participating in newly gained signaling interactions were enriched for their respective stage-specific modules across three contrasts: Pregnancy at E16.5 (odds ratio 17.4, q = 4.7 x 10 ^-18^), Early repair at PPD1 (13.4, 1.6 x10^-20^), and Late repair at PPD7 (7.6, 1.8 x10^-11^). At PPD30, this pattern shifts showing that genes involved in gained interactions demonstrate highest enrichment for the Pregnancy module (11.3), whereas New Homeostasis shows lower relative enrichment (5.4, q = 3.6 x10^-8^). This indicates that the intercellular communication profile at one month postpartum shares structural features with the gestational signaling configuration.

### The tissue settles at a novel homeostatic state

While individual tissue parameters resolve on distinct timelines, the divergence among these organizational levels represents a key finding of this study. Quantifying residual displacement at PPD30 relative to maximal displacement across all stages reveals an ordering: gross anatomical length retains 4% displacement, lineage identity retains 51%, transcriptional state retains 53%, intercellular signaling retains 66%, compartment organization retains 82%, and overall cellular composition retains 88% (Fig. S5). Higher-level morphological metrics thus demonstrate near-complete recovery, whereas molecular, compositional, and signaling architectures remain substantially altered at one month postpartum.

Multiple analytical lines of evidence indicate that the PPD30 state is a distinct homeostatic endpoint and not an incomplete transitional repair phase. First, re-anchoring the analysis to PPD1 demonstrates that PPD30 has diverged from the acute injury state to an extent comparable to the non-pregant control baseline (0.78 versus 0.77 for cellular identity, and 0.61 versus 0.53 for signalling; Fig. S5, proceeding along a developmental trajectory distinct from the pre-pregnancy state. Second, the intercellular interaction network exhibits lower similarity to the non-pregnant control baseline at PPD30 than at PPD7, indicating progressive divergence. Third, all four compartments show concordant directional responses at PPD30 (r = 0.60), correlated with the gestational response direction (r = 0.28), a pattern consistent with coordinated tissue reconfiguration. We designate the New Homeostasis program as the transcriptional signature of this endpoint where 789 genes attaining peak expression at PPD30, concordantly expressed across all four compartments and in 13 of 16 cell types, and enriched for muscle adhesion and conduction, intermediate filament organization, and negative regulation of cytokine signaling.

## Discussion

Here, we provide a whole-uterus single-cell atlas spanning non-pregnant, late pregnancy, parturition, postpartum repair, and the establishment of a new tissue state. By profiling 698,631 cells across five stages, we resolve how epithelial, stromal, endothelial, and immune compartments collectively navigate one of the most extensive physiological remodeling events experienced by an adult organ. The central finding is that postpartum recovery is not a return to the pre-pregnancy state. Although uterine dimensions and epithelial continuity are restored, cellular composition, transcriptional programs, and inferred intercellular communication remain displaced from the non-pregnant configuration. Instead, the uterus establishes a new postpartum transcriptional homeostatic state shaped by the preceding pregnancy-to-repair trajectory.

Four key observations distinguish the PPD30 configuration from a residual acute-injury state. First, the PPD30 state is substantially displaced from the acute PPD1 configuration, indicating progression beyond the immediate injury response. Second, the inferred communication network does not progressively converge on the non-pregnant network but undergoes secondary divergence as postpartum recovery advances. Third, the four major tissue compartments exhibit coordinated directional responses at PPD30 despite executing the preceding transcriptional programs with different timing and amplitude. Finally, the New Homeostasis program peaks at PPD30 across all four compartments and most cell types. Together, these results define the postpartum state not by the persistence of acute damage, but by the coordinated establishment of a transcriptional configuration distinct from both active repair and non-pregnant homeostasis.

The epithelium provides the clearest example of this reconfiguration. It undergoes profound depletion during early repair and is structurally restored by PPD30, yet at that stage it also displays the strongest activation of the New Homeostasis program and the greatest transcriptional divergence from the non-pregnant epithelial state. Its trajectory is therefore not one of simple depletion followed by molecular reinstatement. Instead, epithelial reconstruction incorporates features acquired during pregnancy and repair into the subsequent homeostatic state. In this sense, the restored epithelium is structurally continuous with its pre-pregnancy counterpart but transcriptionally informed by the trajectory through which it was rebuilt.

The persistence of pregnancy-and repair-associated features raises the possibility that the postpartum uterus, specifically the epithelial compartment retains a molecular memory of its previous physiological state. A precedent comes from the skin, where epithelial stem cells exposed to acute inflammation retain chromatin accessibility at selected stress-response loci after inflammation resolves (56). These poised regions enable more rapid transcriptional activation and accelerated barrier restoration following a subsequent injury, demonstrating that epithelial stem cells can encode prior inflammatory experience independently of continued immune-cell input. This adaptation may be beneficial by accelerating future repair, but heightened responsiveness could also increase susceptibility to chronic inflammatory or hyperproliferative disease. Related studies demonstrate that epithelial memory also operates in other systems, including the airway epithelium, where basal cells retain an altered differentiation program after allergic inflammation resolves (57), and the intestine, where mucosal macrophages drive a fetal-like regenerative program in the injured epithelium (81, 82). Postpartum mammary involution offers a parallel within a reproductive organ, the post-lactational gland remodelling through a wound-healing-like, immune-rich program with lasting consequences for later disease risk (54, 55).

Our findings extend this conceptual framework from pathological inflammation to a programmed physiological remodelling event. Pregnancy differs fundamentally from colitis or experimentally induced skin inflammation because it is a recurrent physiological challenge that the uterus is adapted to withstand. Nevertheless, the persistence and integration of pregnancy-and repair-associated transcriptional features after structural reconstruction suggest that physiological experience can also leave a molecular imprint on resident tissue cells. Our data demonstrate a transcriptional imprint but do not establish epigenetic memory. Further studies on persistent chromatin accessibility, histone modifications, DNA methylation or another durable regulatory mechanism, together with a modified response to a subsequent pregnancy or injury, will be required to address how the new homeostatic state is encoded and maintained, and what its functional consequences are. The present atlas identifies the compartments, cell types, and transcriptional programs in which such memory may reside and provides a foundation for determining whether the postpartum state is molecularly encoded, functionally consequential, or both.

Intercellular communication is also reorganized as part of this postpartum state. Total signalling does not decline as repair resolves. Sender and receiver roles shift across stages, signalling load remains elevated, and network composition becomes increasingly distinct from the non-pregnant configuration. At PPD30, epithelial self-preference exceeds its non-pregnant level, stromal-to-epithelial preference remains positive, and immune-to-epithelial preference remains suppressed. The structurally restored epithelium therefore occupies a different relational position within the tissue from its pre-pregnancy counterpart. Postpartum homeostasis may consequently be encoded not only within individual cell states but also across a reconfigured multicellular network. Among the ligands through which the epithelium signals outward, Spp1 is one of the highest-share contributors. We previously identified Spp1 as a macrophage-derived mediator of regeneration in the intestine (81), whether the postpartum epithelium deploys Spp1 in a comparable regenerative capacity should be further investigated.

Beyond its conceptual conclusions, this study establishes a reference atlas of 698,631 cells across five parturition-anchored stages, 16 cell types, and four major tissue compartments. The resource includes cell-level annotations, replicate-resolved composition, five temporal program gene sets with per-contrast statistics, and stage-resolved ligand-receptor interactomes, made available through bulk downloads and an interactive gene-by-cell-type interface. The dataset therefore allows the community to move beyond the global trajectories described here and interrogate when, where, and in which cellular context individual genes, pathways, and candidate signalling relationships operate during pregnancy and postpartum recovery. Whole-organ profiling is particularly valuable because pregnancy remodelling and repair are coordinated multicellular processes that cannot be reconstructed from epithelial, stromal, vascular, or immune datasets in isolation.

Together, the resource and the biological findings establish pregnancy as a powerful model for understanding how adult organs encode physiological history. The uterus restores its dimensions and epithelial continuity, but it does not recreate the molecular configuration that preceded pregnancy. Instead, its major cellular compartments integrate features acquired during pregnancy and repair and collectively establish a new postpartum transcriptional homeostatic state.

## Materials and Methods

### Animals and tissue collection

All in vivo experiments involving mice are adhered to ethical guidelines and regulations and were approved by the Catalonia National Animal Ethics Committees (DMAH12760). Female C57BL/6 mice were obtained from Charles River and housed in the IDIBELL animal facility. Animals were 8 to 16 weeks of age at collection. Females were mated and pregnancies timed to embryonic day 0.5 on the day a copulatory plug was observed. Uteri were collected from non-pregnant control adult females, at embryonic day 16.5, and at postpartum days 1, 7 and 30, with the day of parturition designated postpartum day 0. Three to six independent animals per stage were used for both single-cell profiling and histology. Anatomical and transcriptional data are paired within each animal wherein per animal, one uterine horn was used for histology and one horn was dissociated for single-cell profiling. Horn length was measured using a ruler per uterine horn. For the Non-pregnant control, the estrous cycle stage (Estrus) was confirmed by vaginal cytology, scored on the relative abundance of cornified epithelial cells and leukocytes independently by two researchers as an approximate assignment (58).

### Immunofluorescence

Uterine horns were fixed in 10% neutral-buffered formalin (Sigma-Aldrich Cat # HT501128) overnight at 4 °C and embedded as FFPE blocks. Sections were cut at 5 µm on a Leica RM2255 microtome, dewaxed and subjected to antigen retrieval in citrate buffer pH 6 (Dako/Agilent, Cat # S2369) for 20 min at 95 °C followed by 20 min at room temperature. Sections were permeabilised in 0.1% Triton-X-100 (Life Technologies Cat # 28314) for 30 min and blocked in 0.1% Bovine Serum Albumin (Capricorn Scientific Cat # BSA-1S, Lot No: CP24-7442) in PBS for 1 h. Sections were stained with anti-EpCAM (mouse IgG1, Chemicon/Merck Millipore, Cat # CBL172, 1:50) and anti-FOXA2/HNF3B (rabbit, clone D56D6, Cell Signaling Technology Cat #8186S, 1:100) primary antibodies, followed by donkey anti-mouse IgG Alexa Fluor 647 (Jackson ImmunoResearch, Cat # 715-605-151) and donkey anti-rabbit IgG Alexa Fluor 488 (Jackson ImmunoResearch, Cat # 711-545-152), both at 1:200. Sections were mounted in Fluoromount-G with DAPI (Life Technologies, Cat # 00-4959-52) and imaged on a Zeiss LSM 980 confocal microscope (Carl Zeiss) with a Plan-Apochromat 10x/0.45 M27 objective, acquiring three channels (DAPI, Alexa Fluor 488 and Alexa Fluor 647).

### Tissue dissociation, capture and sequencing

Tissue was minced until the suspension could be drawn through a P1000 pipette tip, then digested in Collagenase type II (Sigma-Aldrich, C9263) at 0.5 mg/mL in HBSS (Sigma-Aldrich Cat # H6648, Lot: RNBM4565) for 45 min at 37 °C with trituration every 15 min. The suspension was filtered sequentially through 100 µm and 70 µm strainers (Pluristrainer Select Cat # V-PM19-2024-01 and 5087560003) and washed in 0.5% bovine serum albumin in PBS. Erythrocytes were removed by 3 min incubation in RBC lysis buffer (BioLegend Cat # 420301, Lot: B319740). Viability was assessed by 7-aminoactinomycin D discrimination (7-AAD; Sigma-Aldrich, CAS 7240-37-1) during sorting on a Beckman Coulter CytoFLEX SRT cell sorter. Digestion time was selected from a 30, 45 and 60 min time-course acquired on a Beckman Coulter CytoFLEX LX analyser by 7-AAD viability and by flow cytometry for EpCAM-BV510 (BioLegend, 118231 clone G8.8,), PDGFRa/CD140a-PE-CF594 (BD Biosciences, 562775,), CD45-FITC ( BioLegend, Cat # 103108, clone 30-F11) and CD31-PE/Cy7 (BioLegend, Cat # 102417, clone 390) to compare per-channel marker recovery across digestion times (Fig. S1a–f). Single-cell suspensions were processed with the 10x Genomics Chromium GEM-X Flex probe-based fixed-RNA assay, v2 Multiplex workflow (60), according to the manufacturer’s instructions, yielding three libraries per stage and 15 libraries in total, of which 14 passed quality control and two were retained at control (Table S1). Libraries were prepared at the Single-Cell Unit of the Josep Carreras Leukaemia Research Institute and sequenced on Illumina NovaSeq. Library quality control is reported in Fig. S1h–l.

### Single-cell data quality control and Seurat object generation

Reads were aligned and quantified with Cell Ranger v10.0 against mouse reference probe set v2.0 (61). The CONTROL sample comprised four animals pooled into two retained libraries, a third control library having yielded 36 cells and failed quality control was not included in the downstream analyses. Per-library filtered count matrices were loaded into Seurat v5.2.1 under R 4.3.3 (62). During object construction, genes detected in fewer than three cells and cells with fewer than 200 detected genes were removed. The mitochondrial fraction per cell was computed from genes matching mt-or MT-. Cells were retained if they had at least 200 detected genes and at least the 0.5th-percentile number of genes, fewer than the 99.5th-percentile total unique molecular identifiers, and no more than 20% mitochondrial content; genes detected in more than two cells were retained. A per-library count-feature regression filter additionally removed cells falling more than 0.4 residual units below the per-library fit of log(nFeature) on log(nCount). log(nCount) fit. Doublets were identified per sequencing unit with scDblFinder v1.16.0 and removed, retaining only cells classified as singlets (63). A total of 698,631 cells passed quality control, distributed as 127,161 (CONTROL), 17,688 (E16.5), 194,125 (PPD1), 338,578 (PPD7) and 21,079 (PPD30). Mean unique molecular identifiers per cell varied substantially across stages (CONTROL 2,279; PPD7 2,460; PPD1 2,881; PPD30 6,076; E16.5 8,285), such that sequencing depth is not independent of stage. Analyses relating transcriptional distance to technical covariates were therefore conditioned on stage.

### Integration, clustering and annotation

Counts were library-size normalized (LogNormalize, scale factor 10,000) and log-transformed, 2,000 highly variable genes were selected by variance-stabilizing transformation, the data were scaled, and 30 principal components were retained. Batch effects were corrected with Harmony over biological replicate (64). A nearest-neighbour graph was constructed on the first 30 Harmony dimensions with k = 20 and clustered by the Louvain algorithm at resolution 0.2, yielding 22 clusters (65). Embeddings were computed with UMAP on the same 30 Harmony dimensions with n.neighbors = 30 and min.dist = 0.3 (66).

Cell types were assigned by collapsing clusters using curated marker gene sets drawn from Tabula Muris Senis and CZ CELLxGENE (67,68) (Fig. 2d and Fig. S2b), and the resulting annotation was cross-validated against the Mouse Cell Atlas uterus reference by per-type correlation (69). Annotation support, resolution sensitivity across 14, 22 and 34 clusters, and the Harmony batch diagnostic are provided in Fig. S2. This yielded 16 cell types. Five types each merge more than one Louvain cluster: Fibroblast (clusters 0, 1 and 18), Stromal (remodeling) (3 and 13), Dendritic cell (19 and 21), Macrophage (2 and 15) and Smooth muscle (8 and 20); the remaining eleven types are single clusters.

Types were grouped into four compartments as follows. Epithelial comprises Epithelial. Endothelial comprises Vascular endothelial and Lymphatic endothelial. Immune comprises Macrophage, Macrophage/APC, Dendritic cell, Dendritic cell (cDC1), T cell and NK cell. Stromal is defined as the residual class and comprises Fibroblast, Fibroblast (Igf1+), Stromal (remodeling), Proliferating stromal, Pericyte, Mesothelial and Smooth muscle. This construction guarantees that no cell type is omitted from the compartment analysis.

### Cell-type composition and Marker expression matrix

Cells per replicate and cell type, and the corresponding within-replicate percentages, were tabulated for all retained libraries. The excluded control library is omitted from every replicate-level statistic while its cells are retained in per-cell projections. Because composition is measured on dissociated cells and sequencing depth covaries with stage, the analysis was repeated after subsampling to 15,000 cells per stage in a single draw at seed 0, which reproduced the direction of every compositional change. Absolute per-replicate composition and the subsampled reproduction are given in Table S2.Per-cluster expression sums were aggregated to cell type and divided by the number of cells of that type, five of the named types comprising two clusters and one comprising three. The resulting matrix was standardized gene-wise across the 16 types, such that the panel reports which cell type expresses each marker most strongly rather than which marker is most abundant within a type. The presented matrix comprises 139 genes.

### Transcriptional module definition and scoring

Modules were derived from whole-tissue pseudobulk differential expression against the non-pregnant control program, with criteria that differ by module and are stated individually. Homeostasis (678 genes) comprises genes higher in CONTROL in at least three of the four control-anchored contrasts. Pregnancy (217), Early repair (985) and Late repair (745) comprise genes elevated relative to CONTROL at E16.5, PPD1 and PPD7 respectively, at q < 0.05 and log2FC ≥ 1. New Homeostasis (789) comprises of genes elevated relative to CONTROL and additionally attain their maximum at PPD30. Homeostasis is defined by consistency across contrasts whereas three modules are defined by a single-contrast threshold. Modules are not mutually exclusive by construction, and their pairwise overlaps are reported in Table S3 part B. Module names denote the stage of maximal expression Gene membership with per-gene log2FC and q value at all four contrasts is provided in Table S3.

### Differential expression and Gene Ontology enrichment

Counts were summed within replicate and cell type at a floor of 50 cells per profile, normalized to counts per million, log-transformed, and tested against the non-pregnant control by empirical-Bayes variance-moderated linear modelling with precision weights (limma-voom) (71,72), with genes declared significant at q < 0.05 and log2FC ≥ after Benjamini-Hochberg correction (73). Aggregation to pseudobulk profiles was doen to avoid inflated False Discovery Rates that follow from treating cells as independent replicates (45–48). Per-compartment persistence, the FDR ceiling and multi-stage comparisons as in Fig. S3. Program gene sets were tested for over-representation of Gene Ontology biological-process terms (74) with Enrichr (75) against the GO_Biological_Process_2023 library and its genome background, with Benjamini–Hochberg correction across terms within each program. The six terms of lowest adjusted P are displayed per program in Fig. 4g, irrespective of whether they reach q < 0.05, and complete results are provided in Table S5.

### Intercellular communication profiling

Ligand-receptor interactions were inferred per stage on the 16-type annotation with LIANA v1.8.1 using the mouse consensus ligand-receptor resource and CellPhoneDB scoring (49, 52,53, 76–78), including all cell types represented by at least 10 cells. Exactly one combination was excluded on this criterion, Fibroblast (Igf1+) at E16.5 with 8 cells, and the remaining 79 stage-by-type combinations were tested. The full coverage log is provided in Fig. S4b, and the per-stage interactions in Table S4. Interactions were inferred from cells pooled within stage. Consequently no replicate-level estimate of communication is available, no confidence interval is placed on signalling load, and interactions represent co-expression-based inference rather than direct measurement (50).

### Cross-compartment coordination and Layer retention analysis

Four-compartment pseudobulk profiles were normalized to counts per million prior to filtering.Three thousand highly variable genes were selected and per-stage log2 fold changes against CONTROL were computed per compartment. Coordination is reported as the mean pairwise correlation between compartment response vectors within stage, the correlation between consensus response vectors across stages, and a null distribution obtained by permuting gene labels. Retention at PPD30 was computed per organizational layer as the fraction of that layer’s maximum displacement from CONTROL across stages that remains at PPD30, using the layer’s native distance measure: absolute difference for horn length, correlation distance on composition vectors for composition, correlation distance on pseudobulk log2FC vectors for transcriptional state and compartment organization, and Jaccard distance on the interaction edge set for signalling. The same quantities were recomputed with PPD1 as the anchor in place of CONTROL to distinguish an unfinished trajectory from a completed one with a different destination. Both re-anchored analyses are shown in Fig. S5.

### Statistics

The biological replicate is one animal per sequencing library at E16.5, PPD1, PPD7 and PPD30; For Control, four animals were pooled into two retained libraries. For horn length the replicate is the horn. Tests and exact sample sizes are stated in each figure legend. Multiple comparisons were corrected as stated per analysis, and all software packages, versions and random seeds are specified in the analysis repository (see Data and code availability).

## Acknowledgments

We thank the IDIBELL Animal Facility for animal husbandry and colony management, the IDIBELL Histology, Flow Cytometry and Optical Microscopy units for tissue processing, cell sorting and imaging support, and the Single-Cell Unit of the Josep Carreras Leukaemia Research Institute for library preparation and sequencing. We acknowledge the Novo Nordisk Foundation Center for Stem Cell Medicine (reNEW) and P-CMR[C] server and computing cluster facilities. We thank members of the Cell Plasticity and Regeneration Guiu Lab for feedback and discussion.

## Author contributions

Conceptualization: JMT, JG. Experiments: JMT, OW, BA, JG. Analysis: JMT, LM. Visualization and Plotting: JMT. Writing – first draft: JMT. Writing, review and editing: all authors. Supervision: JG. Funding acquisition: JMT, JG.

## Data, code, and materials availability

All data needed to evaluate the conclusions in the paper are present in the paper and the Supplementary Materials. Raw sequencing data are deposited at Gene Expression Omnibus. Analysis code is available at github.com/jmyteves/sc_endom_regeneration, currently a private repository, which will be made public upon acceptance.

## Supplementary Materials

This PDF file includes Figs. S1 to S5, the supplementary figure legends and the table descriptions. ALL_Tables.xlsx includes Tables S1 to S6.

