## Supplemental Figures S1-S5 for "Mapping Uterine Remodeling from Pregnancy to a New Homeostatic State"

Supplementary Materials for
**Systematic mapping of the endometrium towards new homeostasis**
Joji Marie Teves et al.

**This PDF file includes:**

Figs. S1 to S5
Tables S1 to S6 (descriptions; data supplied as a separate workbook)

**Other Supplementary Materials for this manuscript include the following:**
Data S1 (ALL_Tables.xlsx) — Tables S1 to S6 as a single workbook, one sheet per table.

**Fig. S1. Dissociation optimization and library quality control.** (a-c) Representative tube per digestion time, gated on singlets, (a) 30 min, (b) 45 min (c) and 60 min. (d) Viability, 7-AAD-negative as a percentage of singlets (81.0, 56.4 and 60.0 percent). (e) Live-cell yield per mg of input tissue (5.9, 12.7 and 3.0). (f) Per-channel positive fraction of live cells, with 45 min marked as the condition taken forward; bars are means of replicate tubes, error bars SD and points individual tubes. (g) Library medians on the count-feature relationship, log median genes against log median unique molecular identifiers per cell; the fit has slope 0.654, r = 0.970 and R2 = 0.942 across the 14 retained libraries, and the band marks the 0.4-residual exclusion boundary below the fit, with E16.5-3 carrying the largest residual at -0.27 and retained. (h) Cells passing quality control per library, all fifteen prepared libraries shown, with the excluded CTRL-3 library annotated. (i) Cells recovered per stage after quality control: 127k control, 18k E16.5, 194k PPD1, 339k PPD7 and 21k PPD30, totalling 698,595 cells across the 14 retained libraries. (j) Detected genes per cell by stage, medians 1,141, 2,384, 1,569, 1,580 and 2,519, with the 200-gene minimum drawn. (k) Total unique molecular identifiers per cell by stage, medians 1,548, 5,784, 2,500, 2,299 and 4,877. (l) Mitochondrial fraction by stage, medians 4.7, 5.1, 3.6, 2.2 and 4.5 percent, with the 20 percent maximum drawn. n = 3 replicate tubes at 30 and 45 min and n = 2 at 60 min.

**Fig. S2. Annotation validity, batch integration and clustering robustness.** (a) Embeddings before and after Harmony integration, colored by cell type and by stage, showing that within-stage replicates mix after correction while cell-type structure is preserved. (b) Marker specificity dot plot across the annotated types, with dot size giving the fraction of cells in the group expressing each gene and color giving mean expression within the group. (c) Quality-control metrics overlaid on the embedding: log10 unique molecular identifiers, log10 detected genes, percent mitochondrial reads and doublet score. (d) Cells recovered per replicate and within-replicate cell-type composition, grouped and colored by stage. (e) Pseudobulk signature correlation between our control cell types and Mouse Cell Atlas uterus labels, computed on highly variable genes; asterisks mark each row's best-correlating reference label. (f) Compartment-level concordance between our control compartments and the Mouse Cell Atlas uterus compartments, with every diagonal entry exceeding its off-diagonal alternatives (epithelial 0.49, stromal 0.57, muscle 0.61, endothelial 0.59, mesothelial 0.73, immune 0.69).

**Fig. S3. Transcriptional program robustness.** (a) Control-anchored differential expression at each stage, plotted as log2fold change against −log10 adjusted P: control versus E16.5 (1,361 control-high, 222 E16.5-high), versus PPD1 (2,045 and 985), versus PPD7 (1,368 and 745) and versus PPD30 (1,268 and 1,089). (b) Mean gene z score per program across stages, with points distinguished by whether they fall outside or inside the 95 percent interval of a size- and expression-matched random gene-set null. (c) Shared gene counts between the five program sets; set sizes are 678 Homeostasis, 217 Pregnancy, 985 Early repair, 745 Late repair and 789 New Homeostasis, and the largest overlap is 189 genes between Early repair and Late repair. (d)

The same overlaps as Jaccard index, maximum 12.3 percent, establishing that the programs are near-disjoint despite not being mutually exclusive by construction. (e) Program expression resolved by compartment, shown as pseudobulk z-scores per stage for epithelial, stromal, immune and endothelial compartments (20k, 497k, 155k and 26k cells respectively), alongside Gene Ontology Biological Process enrichment for the five programs, with dot size giving genes in term and color giving −log10 adjusted P.

**Fig. S4. Communication inference controls and epithelial ligand-receptor detail.** (a) Inferred interaction totals against cells sequenced per stage, for active ligand-receptor interactions at $P < 0.05$ across 16 cell types (Spearman $\rho = -0.60$, $P = 0.28$), signalling load across the four compartments ($\rho = -0.90$, $P = 0.04$) and total directed edges across all 16 × 16 type pairs ($\rho = -0.60$, $P = 0.28$). (b) Cells available per cell type and stage on a log scale. (c) The fourteen highest-signal ligand-receptor pairs for each of the four principal compartment routes: stromal to epithelial, epithelial to stromal, immune to epithelial and epithelial to immune, with dot size giving total signal and color giving the control share of that pair's signal. (d) Epithelial share of signal resolved to individual ligands, as receiver from all other types and as sender to all other types, with color diverging about equal control and stage contribution.

**Fig. S5. Compartment coordination and anchor sensitivity of the layer analysis.**

(a) Pearson correlation between compartment response vectors at each postpartum stage. (b) Layer retention recomputed under three anchor choices. (c) Mean correlation of each compartment to the other three across stages. (d) Anchor sensitivity per layer, given as the correlation between the control-anchored and re-anchored retention profiles.

**Data S1. (separate file)**

**ALL_Tables.xlsx.** Tables S1 to S6 within a single workbook represented as one sheet per table.

**Table S1.**

Library and sample metrics. Per-library metrics for the 14 libraries retained after quality control, spanning five stages with three biological replicates per stage and two for control. Columns give library identifier, stage, replicate index within stage, cells recovered after quality control, median unique molecular identifiers and median genes per cell, percent mitochondrial reads and percent doublets flagged. Total 698,595 cells.

**Table S2.**

Per-library cell-type composition. Cell-type composition of each of the 14 libraries, 224 rows covering all 16 cell types in every library. Columns give library identifier, stage, cell type, cells of that type, library total, percent within library, the delivered percentage and the absolute difference between the two. The maximum discrepancy is below $1.5 \times 10^{-14}$ percentage points, so the table serves as a reconciliation check between computed and delivered composition.

**Table S3.**

Transcriptional program gene sets and their pairwise overlaps.  Every gene assigned to each of the five programs: Early repair 985, New Homeostasis 789, Late repair 745, Homeostasis 678 and Pregnancy 217.  with log2fold change and Benjamini-Hochberg adjusted q at all four control-anchored contrasts, the stage at which the program peaks, and the rule defining the program. Part B (10 rows) gives the pairwise overlaps between the five sets.

**Table S4.**

Ligand-receptor interactions per stage. 5,760 rows covering 1,694 distinct ligand-receptor pairs across the four control-anchored contrasts (E16.5 1,494, PPD1 1,381, PPD7 1,367, PPD30 1,518). For each pair and contrast the table gives the number of cell-type pairs tested, mean interaction value in control and at stage, mean control share, the mean, median, maximum and minimum shift toward the stage, and counts of cell-type pairs gained, lost or absent in control.

**Table S5.**

Gene Ontology Biological Process enrichment per transcriptional program. All 4,673 Biological Process terms tested against each of the five program gene sets, 12,135 term-program combinations in total. Columns give the term, raw P, Benjamini-Hochberg adjusted P, odds ratio, combined score, overlapping genes, program, overlap size, program set size and gene ratio.

**Table S6.**

Per-compartment pseudobulk temporal modules and enrichment of the New Homeostasis module. Unbiased temporal modules derived independently within each of the four compartments, and the Gene Ontology enrichment of the PPD30-peaking module; source of Fig. 5. Three parts in one sheet, distinguished by the _part column. Part A (6,000 rows) gives every gene-by-module assignment for all four compartments: the 1,500 genes of highest one-way ANOVA F across the five stages of log1p-CPM pseudobulk per compartment, each assigned to one of five k-means modules on stage z scores, with that gene's z in every stage, its own peak stage, and flags for whether that peak is PPD30 and whether it is up at PPD30 only. Part B (20 rows) is the four-compartment by five-module summary giving module size, mean z per stage, peak stage, the peak-minus-runner-up z margin and the k = 5 silhouette. Part C (6,348 rows) is every Gene Ontology biological-process term returned for each compartment's New Homeostasis module, tested against an all-genes genome background with Benjamini-Hochberg correction, with a flag marking the ten terms per compartment drawn in Fig. 5. The PPD30-peaking module is M5 in all four compartments: epithelial 313 genes, stromal 187, immune 223 and endothelial 283. Two caveats apply to cross-compartment comparison. The four enrichment analyses are independent and share no common term axis, and their significance is uneven. Across all terms returned, the number reaching $q < 0.05$ is 10 (epithelial), 16 (immune), 1 (endothelial) and none (stromal, lowest adjusted P 0.146); among the ten terms drawn per compartment in Fig. 5 the corresponding counts are 10, 10, 1 and none, which is what the panel annotations report. Epithelial M5 is a composite rather than a clean PPD30 module, its mean z being nearly tied between PPD1 and PPD30 and only 155 of its 313 genes peaking individually at PPD30. Enrichment here used the GO_Biological_Process_2025 library, whereas the whole-tissue program enrichment in Table S5 used GO_Biological_Process_2023.

### Fig. S1

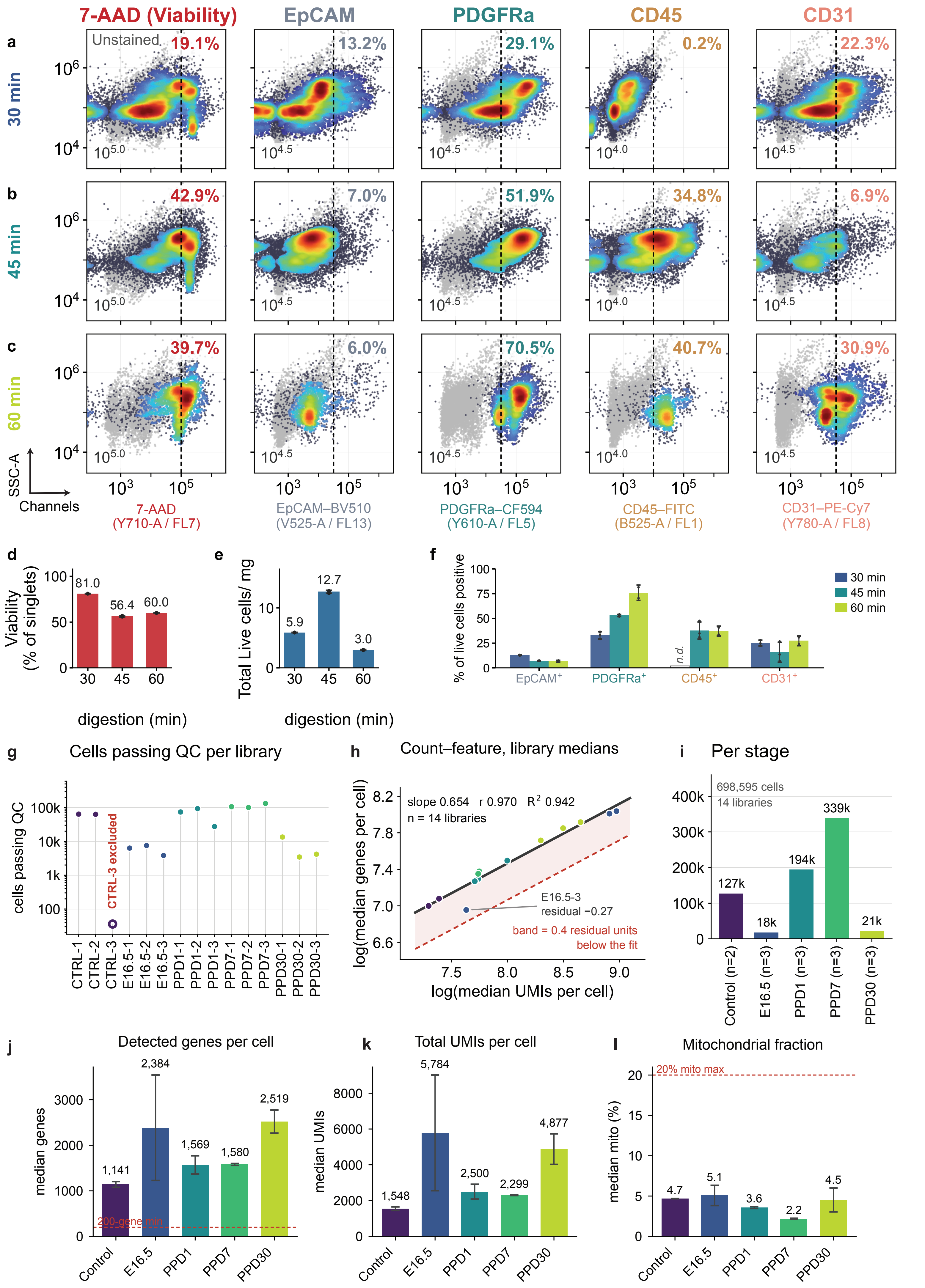
7-AAD (Viability)
EpCAM
PDGFRa
CD45
CD31
a
30 min
Unstained
19.1%
13.2%
29.1%
0.2%
22.3%
b
45 min
42.9%
7.0%
51.9%
34.8%
6.9%
c
60 min
39.7%
6.0%
70.5%
40.7%
30.9%
SSC-A
Channels
7-AAD (Y710-A / FL7)
EpCAM–BV510 (V525-A / FL13)
PDGFRa–CF594 (Y610-A / FL5)
CD45–FITC (B525-A / FL1)
CD31–PE-Cy7 (Y780-A / FL8)
d
Viability (% of singlets)
81.0
56.4
60.0
digestion (min)
e
Total Live cells/ mg
5.9
12.7
3.0
digestion (min)
f
% of live cells positive
30 min
45 min
60 min
n.d.
EpCAM+
PDGFRa+
CD45+
CD31+
g
Cells passing QC per library
cells passing QC
CTRL-3 excluded
CTRL-1
CTRL-2
CTRL-3
E16.5-1
E16.5-2
E16.5-3
PPD1-1
PPD1-2
PPD1-3
PPD7-1
PPD7-2
PPD7-3
PPD30-1
PPD30-2
PPD30-3
h
Count–feature, library medians
slope 0.654 r 0.970 R2 0.942
n = 14 libraries
E16.5-3
residual −0.27
band = 0.4 residual units below the fit
log(median genes per cell)
log(median UMIs per cell)
i
Per stage
698,595 cells
14 libraries
127k
18k
194k
339k
21k
Control (n=2)
E16.5 (n=3)
PPD1 (n=3)
PPD7 (n=3)
PPD30 (n=3)
j
Detected genes per cell
median genes
1,141
2,384
1,569
1,580
2,519
200-gene min
k
Total UMIs per cell
median UMIs
1,548
5,784
2,500
2,299
4,877
l
Mitochondrial fraction
median mito (%)
20% mito max
4.7
5.1
3.6
2.2
4.5
Control
E16.5
PPD1
PPD7
PPD30

Fig. S2

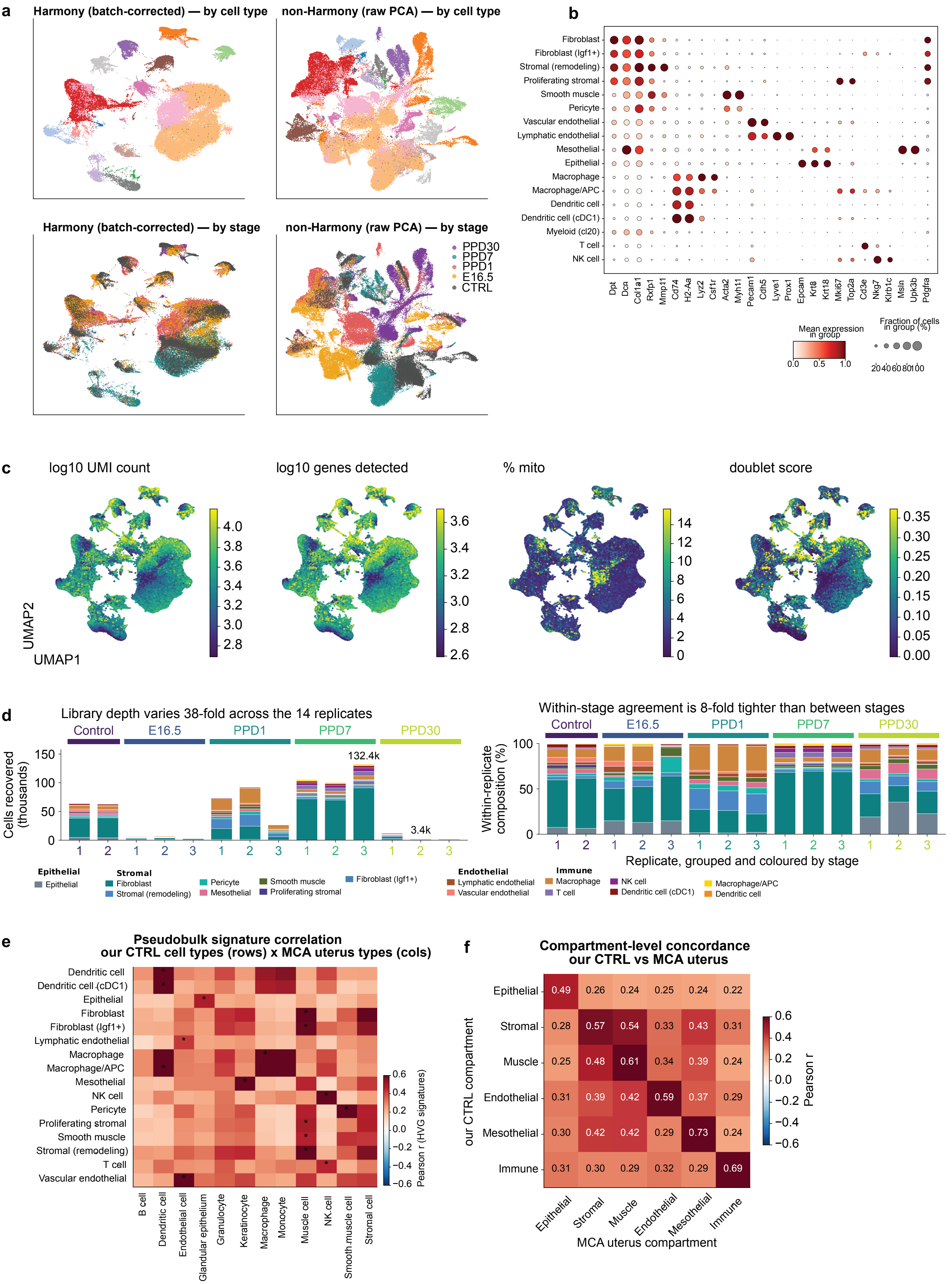
a
Harmony (batch-corrected) — by cell type
non-Harmony (raw PCA) — by cell type
Harmony (batch-corrected) — by stage
non-Harmony (raw PCA) — by stage
PPD30
PPD7
PPD1
E16.5
CTRL
b
Fibroblast
Fibroblast (Igf1+)
Stromal (remodeling)
Proliferating stromal
Smooth muscle
Pericyte
Vascular endothelial
Lymphatic endothelial
Mesothelial
Epithelial
Macrophage
Macrophage/APC
Dendritic cell
Dendritic cell (cDC1)
Myeloid (cl20)
T cell
NK cell
Dpt
Dcn
Col1a1
Rxfp1
Mmp11
Cd74
H2-Aa
Lyz2
Csf1r
Acta2
Myh11
Pecam1
Cdh5
Lyve1
Prox1
Epcam
Krt8
Krt18
Mki67
Top2a
Cd3e
Nkg7
Klrb1c
Msln
Upk3b
Pdgfra
Mean expression in group
0.0
0.5
1.0
Fraction of cells in group (%)
20406080100
c
log10 UMI count
log10 genes detected
% mito
doublet score
UMAP2
UMAP1
d
Library depth varies 38-fold across the 14 replicates
Control
E16.5
PPD1
PPD7
PPD30
Cells recovered (thousands)
132.4k
3.4k
Within-stage agreement is 8-fold tighter than between stages
Within-replicate composition (%)
Replicate, grouped and coloured by stage
Epithelial
Epithelial
Stromal
Fibroblast
Stromal (remodeling)
Pericyte
Mesothelial
Smooth muscle
Proliferating stromal
Fibroblast (Igf1+)
Endothelial
Lymphatic endothelial
Vascular endothelial
Immune
Macrophage
T cell
NK cell
Dendritic cell (cDC1)
Macrophage/APC
Dendritic cell
e
Pseudobulk signature correlation
our CTRL cell types (rows) x MCA uterus types (cols)
Dendritic cell
Dendritic cell (cDC1)
Epithelial
Fibroblast
Fibroblast (Igf1+)
Lymphatic endothelial
Macrophage
Macrophage/APC
Mesothelial
NK cell
Pericyte
Proliferating stromal
Smooth muscle
Stromal (remodeling)
T cell
Vascular endothelial
B cell
Dendritic cell
Endothelial cell
Glandular epithelium
Granulocyte
Keratinocyte
Macrophage
Monocyte
Muscle cell
NK cell
Smooth muscle cell
Stromal cell
Pearson r (HVG signatures)
f
Compartment-level concordance
our CTRL vs MCA uterus
our CTRL compartment
MCA uterus compartment
Pearson r
Epithelial 0.49 0.26 0.24 0.25 0.24 0.22
Stromal 0.28 0.57 0.54 0.33 0.43 0.31
Muscle 0.25 0.48 0.61 0.34 0.39 0.24
Endothelial 0.31 0.39 0.42 0.59 0.37 0.29
Mesothelial 0.30 0.42 0.42 0.29 0.73 0.24
Immune 0.31 0.30 0.29 0.32 0.29 0.69
Epithelial
Stromal
Muscle
Endothelial
Mesothelial
Immune

Fig. S3

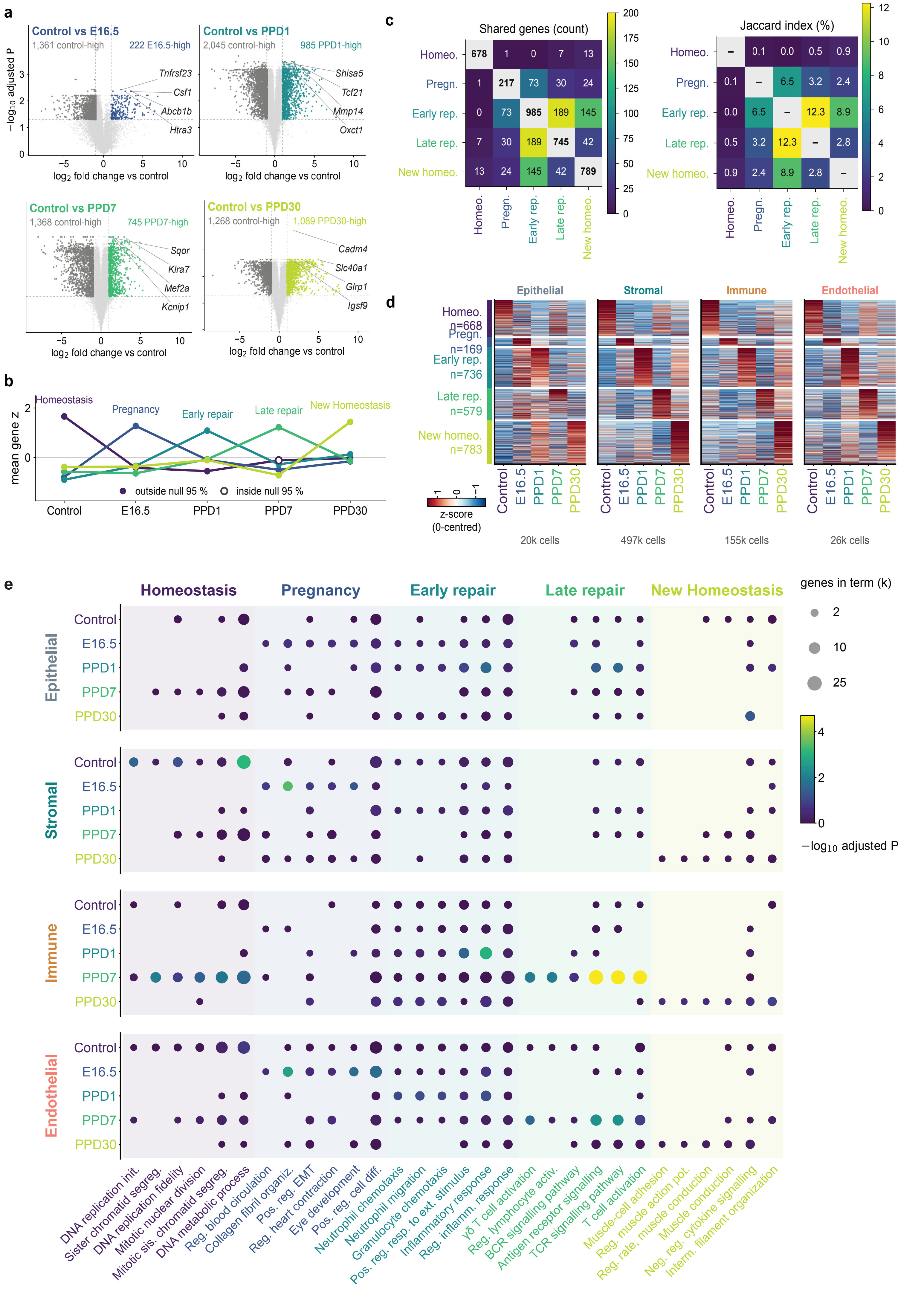

a
Control vs E16.5
1,361 control-high
222 E16.5-high
Tnfrsf23
Csf1
Abcb1b
Htra3
Control vs PPD1
2,045 control-high
985 PPD1-high
Shisa5
Tcf21
Mmp14
Oxct1
Control vs PPD7
1,368 control-high
745 PPD7-high
Sqor
Klra7
Mef2a
Kcnip1
Control vs PPD30
1,268 control-high
1,089 PPD30-high
Cadm4
Slc40a1
Glrp1
Igsf9
−log10 adjusted P
log2 fold change vs control
b
Homeostasis
Pregnancy
Early repair
Late repair
New Homeostasis
mean gene z
outside null 95 %
inside null 95 %
Control
E16.5
PPD1
PPD7
PPD30
c
Shared genes (count)
Homeo. 678 1 0 7 13
Pregn. 1 217 73 30 24
Early rep. 0 73 985 189 145
Late rep. 7 30 189 745 42
New homeo. 13 24 145 42 789
Jaccard index (%)
Homeo. – 0.1 0.0 0.5 0.9
Pregn. 0.1 – 6.5 3.2 2.4
Early rep. 0.0 6.5 – 12.3 8.9
Late rep. 0.5 3.2 12.3 – 2.8
New homeo. 0.9 2.4 8.9 2.8 –
d
Epithelial
Stromal
Immune
Endothelial
Homeo. n=668
Pregn. n=169
Early rep. n=736
Late rep. n=579
New homeo. n=783
z-score (0-centred)
20k cells
497k cells
155k cells
26k cells
e
Homeostasis
Pregnancy
Early repair
Late repair
New Homeostasis
genes in term (k)
2
10
25
−log10 adjusted P
Epithelial
Stromal
Immune
Endothelial
DNA replication init.
Sister chromatid segreg.
DNA replication fidelity
Mitotic nuclear division
Mitotic sis. chromatid segreg.
DNA metabolic process
Reg. blood circulation
Collagen fibril organiz.
Pos. reg. EMT
Reg. heart contraction
Eye development
Pos. reg. cell diff.
Neutrophil chemotaxis
Neutrophil migration
Granulocyte chemotaxis
Pos. reg. resp. to ext. stimulus
Inflammatory response
Reg. inflamm. response
γδ T cell activation
Reg. lymphocyte activ.
BCR signalling pathway
Antigen receptor signalling
TCR signalling pathway
T cell activation
Muscle-cell adhesion
Reg. muscle action pot.
Reg. rate, muscle conduction
Muscle conduction
Neg. reg. cytokine signalling
Interm. filament organization

### Fig. S4

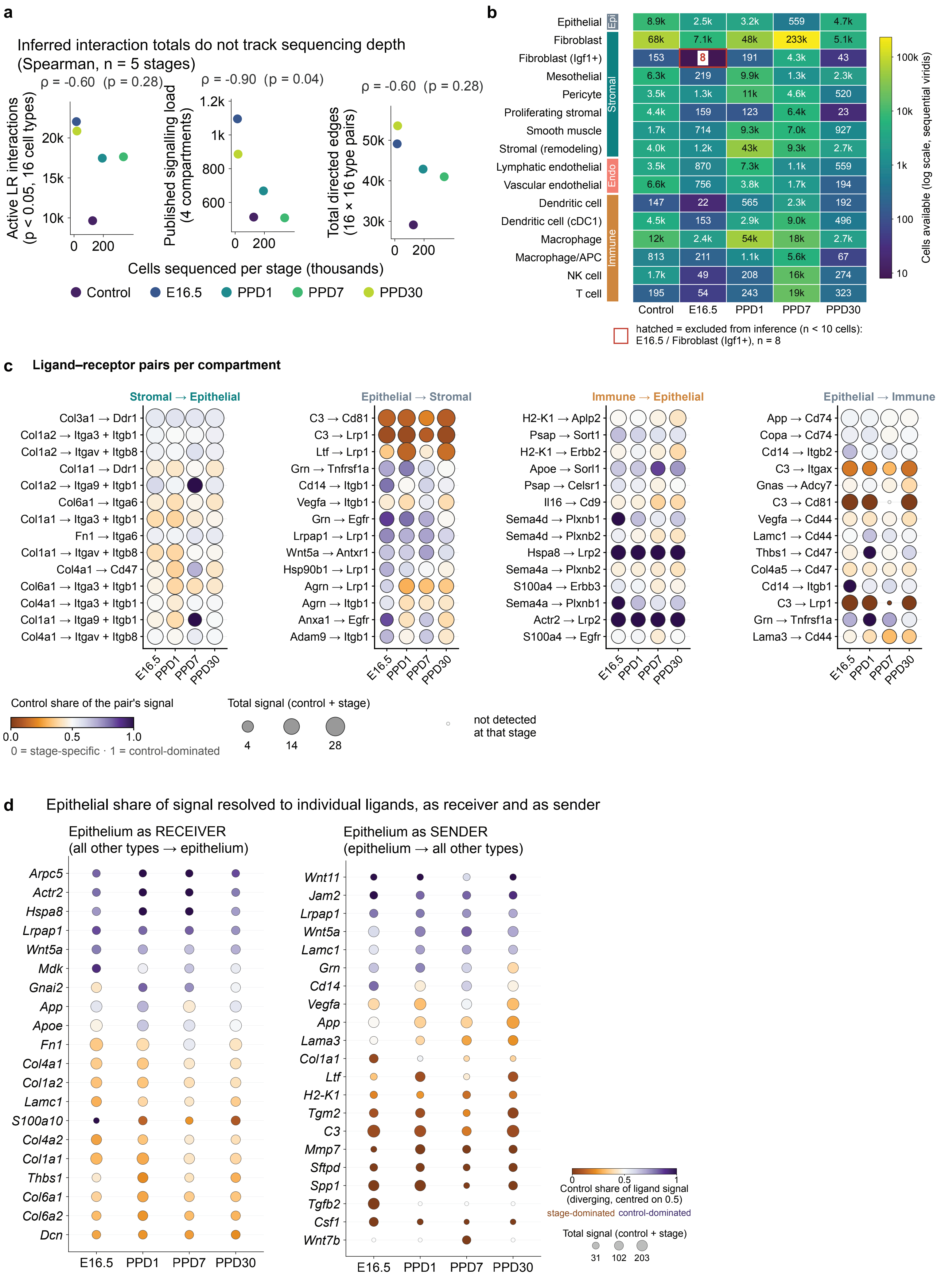

a
Inferred interaction totals do not track sequencing depth
(Spearman, n = 5 stages)
ρ = -0.60 (p = 0.28)
ρ = -0.90 (p = 0.04)
ρ = -0.60 (p = 0.28)
Active LR interactions
(p < 0.05, 16 cell types)
Published signalling load
(4 compartments)
Total directed edges
(16 × 16 type pairs)
Cells sequenced per stage (thousands)
Control
E16.5
PPD1
PPD7
PPD30
b
Epithelial 8.9k 2.5k 3.2k 559 4.7k
Fibroblast 68k 7.1k 48k 233k 5.1k
Fibroblast (Igf1+) 153 8 191 4.3k 43
Mesothelial 6.3k 219 9.9k 1.3k 2.3k
Pericyte 3.5k 1.3k 11k 4.6k 520
Proliferating stromal 4.4k 159 123 6.4k 23
Smooth muscle 1.7k 714 9.3k 7.0k 927
Stromal (remodeling) 4.0k 1.2k 43k 9.3k 2.7k
Lymphatic endothelial 3.5k 870 7.3k 1.1k 559
Vascular endothelial 6.6k 756 3.8k 1.7k 194
Dendritic cell 147 22 565 2.3k 192
Dendritic cell (cDC1) 4.5k 153 2.9k 9.0k 496
Macrophage 12k 2.4k 54k 18k 2.7k
Macrophage/APC 813 211 1.1k 5.6k 67
NK cell 1.7k 49 208 16k 274
T cell 195 54 243 19k 323
Control E16.5 PPD1 PPD7 PPD30
Epi
Stromal
Endo
Immune
Cells available (log scale, sequential viridis)
hatched = excluded from inference (n < 10 cells):
E16.5 / Fibroblast (Igf1+), n = 8
c Ligand–receptor pairs per compartment
Stromal → Epithelial
Col3a1 → Ddr1
Col1a2 → Itga3 + Itgb1
Col1a2 → Itgav + Itgb8
Col1a1 → Ddr1
Col1a2 → Itga9 + Itgb1
Col6a1 → Itga6
Col1a1 → Itga3 + Itgb1
Fn1 → Itga6
Col1a1 → Itgav + Itgb8
Col4a1 → Cd47
Col6a1 → Itga3 + Itgb1
Col4a1 → Itga3 + Itgb1
Col1a1 → Itga9 + Itgb1
Col4a1 → Itgav + Itgb8
Epithelial → Stromal
C3 → Cd81
C3 → Lrp1
Ltf → Lrp1
Grn → Tnfrsf1a
Cd14 → Itgb1
Vegfa → Itgb1
Grn → Egfr
Lrpap1 → Lrp1
Wnt5a → Antxr1
Hsp90b1 → Lrp1
Agrn → Lrp1
Agrn → Itgb1
Anxa1 → Egfr
Adam9 → Itgb1
Immune → Epithelial
H2-K1 → Aplp2
Psap → Sort1
H2-K1 → Erbb2
Apoe → Sorl1
Psap → Celsr1
Il16 → Cd9
Sema4d → Plxnb1
Sema4d → Plxnb2
Hspa8 → Lrp2
Sema4a → Plxnb2
S100a4 → Erbb3
Sema4a → Plxnb1
Actr2 → Lrp2
S100a4 → Egfr
Epithelial → Immune
App → Cd74
Copa → Cd74
Cd14 → Itgb2
C3 → Itgax
Gnas → Adcy7
C3 → Cd81
Vegfa → Cd44
Lamc1 → Cd44
Thbs1 → Cd47
Col4a5 → Cd47
Cd14 → Itgb1
C3 → Lrp1
Grn → Tnfrsf1a
Lama3 → Cd44
E16.5 PPD1 PPD7 PPD30
Control share of the pair's signal
0.0 0.5 1.0
0 = stage-specific · 1 = control-dominated
Total signal (control + stage)
4 14 28
not detected
at that stage
d Epithelial share of signal resolved to individual ligands, as receiver and as sender
Epithelium as RECEIVER
(all other types → epithelium)
Arpc5
Actr2
Hspa8
Lrpap1
Wnt5a
Mdk
Gnai2
App
Apoe
Fn1
Col4a1
Col1a2
Lamc1
S100a10
Col4a2
Col1a1
Thbs1
Col6a1
Col6a2
Dcn
E16.5 PPD1 PPD7 PPD30
Epithelium as SENDER
(epithelium → all other types)
Wnt11
Jam2
Lrpap1
Wnt5a
Lamc1
Grn
Cd14
Vegfa
App
Lama3
Col1a1
Ltf
H2-K1
Tgm2
C3
Mmp7
Sftpd
Spp1
Tgfb2
Csf1
Wnt7b
E16.5 PPD1 PPD7 PPD30
0 0.5 1
Control share of ligand signal
(diverging, centred on 0.5)
stage-dominated control-dominated
Total signal (control + stage)
31 102 203

### Fig. S5

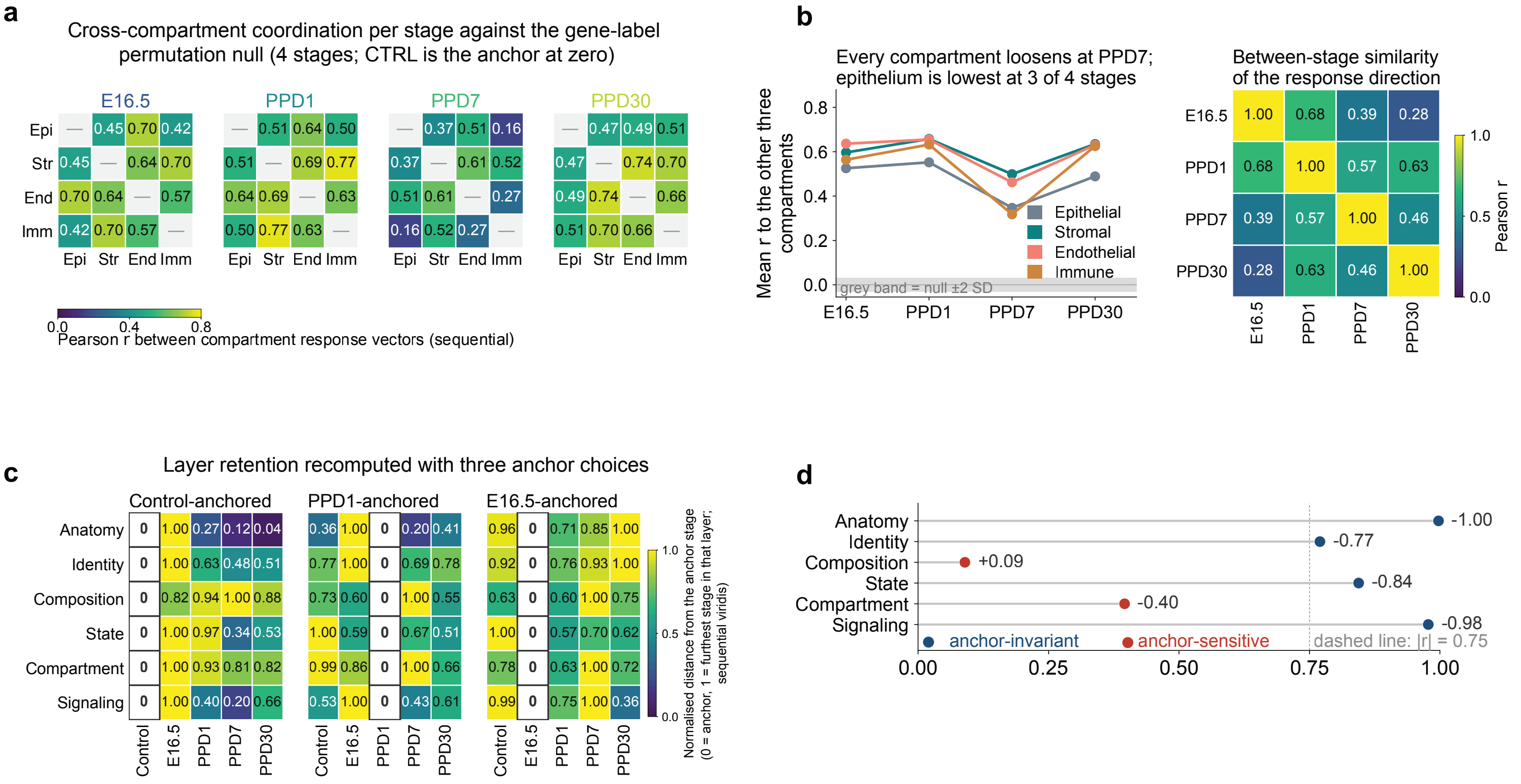
a
Cross-compartment coordination per stage against the gene-label permutation null (4 stages; CTRL is the anchor at zero)
E16.5
PPD1
PPD7
PPD30
Epi Str End Imm
Pearson r between compartment response vectors (sequential)
b
Every compartment loosens at PPD7; epithelium is lowest at 3 of 4 stages
Mean r to the other three compartments
grey band = null ±2 SD
Epithelial
Stromal
Endothelial
Immune
E16.5 PPD1 PPD7 PPD30
Between-stage similarity of the response direction
Pearson r
c
Layer retention recomputed with three anchor choices
Control-anchored
PPD1-anchored
E16.5-anchored
Anatomy
Identity
Composition
State
Compartment
Signaling
Control E16.5 PPD1 PPD7 PPD30
Normalised distance from the anchor stage (0 = anchor, 1 = furthest stage in that layer; sequential viridis)
d
Anatomy -1.00
Identity -0.77
Composition +0.09
State -0.84
Compartment -0.40
Signaling -0.98
anchor-invariant
anchor-sensitive
dashed line: |r| = 0.75
